# Lack of specificity of planarian progenitor responses to injury in regeneration

**DOI:** 10.64898/2026.02.19.706786

**Authors:** Cecilia E. Pellegrini, Peter W. Reddien

## Abstract

Regeneration is the process by which organisms replace lost body parts. How cell-type production is tailored to match the identity of missing tissues is a central problem of regeneration. Here, we investigated the specificity of planarian stem-cell responses to the identity of missing tissues following injury. Proximal injury not affecting the mature tissue nonetheless drives increased cell incorporation into brain neurons, ventral nerve cords, pharynx muscle and neurons. Following direct injury, peripheral neurons show a spatially imprecise, generic amplification of incorporation relative to the injury; body-wall muscle incorporation was amplified, with decreased incorporation at wound-distal sites in favor of increased incorporation wound-proximally. By contrast, essentially no stem-cell division contributes to initial epidermal regeneration, instead post-mitotic progenitors supply the wound. Amplification of epidermal incorporation following injury does occur weeks after injury, including to uninjured regions. These results indicate that the identity of the missing mature tissue is not required in determining the stem-cell response to injury. We suggest that planarian regeneration specificity involves a combination of ongoing cell turnover, wound-associated amplification of stem-cells, and spatially broad neoblast specification zones.

## Introduction

At some stage of their existence most animals will be challenged to repair injured tissues. Many organisms are only capable of local repair, others can regenerate entire regions of body axes (Brockes and Kumar, 2008; Poss, 2010; Srivastava, 2021). In some contexts, regeneration involves constitutively active stem-cells (Rink, 2018; Tanaka and Reddien, 2011; Wagner et al., 2011) that respond to injury by accelerating new-tissue production to replace those that are lost. How organisms are able to interpret tissue absence and carry out its specific replacement remains poorly understood.

Planarians regenerate from a myriad of injuries, being capable of replacing essentially any organ in a matter of days. Planarian regeneration, like that of many regenerative organisms, involves the formation of an outgrowth called a blastema, where differentiation of many new tissues occurs (Reddien, 2018). The planarian body plan is relatively simple, yet highly conserved, including: a nervous system, intestine, musculature, epidermis, kidney-like protonephridia, gland cells, gonads, germline, and phagocytic cells (Hyman, 1951; Reddien and Sánchez Alvarado, 2004).

Underlying planarian homeostatic tissue maintenance and regeneration is a population of stem-cells called neoblasts (Baguñà et al., 1989; Reddien and Sánchez Alvarado, 2004; Rink, 2018; Wagner et al., 2011; Zhu and Pearson, 2016). Neoblasts are the only dividing somatic cells of the animal, are distributed throughout the parenchyma (Reddien and Sánchez Alvarado, 2004), are pluripotent at the population and single-cell levels (Baguñà et al., 1989; Wagner et al., 2011), and express specific markers such as the Piwi-family gene *smedwi-1* (Guo et al., 2006; Reddien et al., 2005). Many neoblasts are considered specialized (fate-specified), expressing Fate-Specific Transcription Factors (FSTFs) (Raz et al., 2021; Scimone et al., 2014; van Wolfswinkel et al., 2014). Specialized neoblasts give rise to specific tissues, and many of these transcription factors are required for homeostatic and regenerative production of particular cell-types (Lapan and Reddien, 2011; Reddien, 2022; Scimone et al., 2011; Zhu and Pearson, 2016). Planarian tissues undergo constant turnover – balancing neoblast-derived cell production and cell death (Pellettieri and Sánchez Alvarado, 2007; Pellettieri et al., 2010; Reddien, 2024). When challenged by injury, neoblast proliferation rapidly increases broadly across the body1-6 hrs post-injury (hpi), as part of the “Generic Wound Response” (GWR) (Baguñà, 1976; Saló and Baguñà, 1984; Wenemoser and Reddien, 2010). As the name indicates, this response occurs for a wide array of injuries, regardless of tissue loss. When substantial tissue loss occurs, a second set of responses follows, collectively known as the “Missing Tissue Response” (MTR). The MTR includes a 48 hpi peak of wound-proximal sustained proliferation, sustained wound-induced gene expression ∼24 hpi, and a body-wide increase in apoptosis associated with tissue re-scaling (Pellettieri et al., 2010; Wenemoser and Reddien, 2010; Wenemoser et al., 2012; Wurtzel et al., 2015).

Planarian tissue regeneration and maintenance is guided by constitutively present adult positional information (Reddien, 2011) involving positional control genes (PCGs) that are regionally and constitutively expressed along body axes, typically in muscle (Cebrià, 2016; Lander and Petersen, 2016; Reddien, 2018; Scimone et al., 2016; Witchley et al., 2013). Neoblast fate-specification occurs in a regional, although spatially coarse, manner; likely influenced by PCG expression domains (Park et al., 2023; Reddien, 2018; Witchley et al., 2013; Wurtzel et al., 2017).

Varying planarian injuries confer distinct regenerative problems. How does the system ‘know’ what tissues are missing? Some studies posit that progenitor responses to injury are guided by the presence/absence of their target tissues (surveillance) (Fig. 1A) (Adler et al., 2014; Bohr et al., 2021; Vu et al., 2015). Another study suggested that the identity of the injured differentiated tissue (i.e., the planarian eye) carries little relevance for the specificity of progenitor responses (LoCascio et al., 2017). In this model, regeneration was referred to as “target-blind”, a situation where mature tissue absence is not sufficient to increase its own progenitor production; when wounding elicits a proliferative response, the types of progenitors amplified is guided by the PCG-defined location of the wound (LoCascio et al., 2017). These differing models highlight the fundamental nature of the question: what is the relationship between lost-tissue identity and stem cell responses?

**Figure 1:**
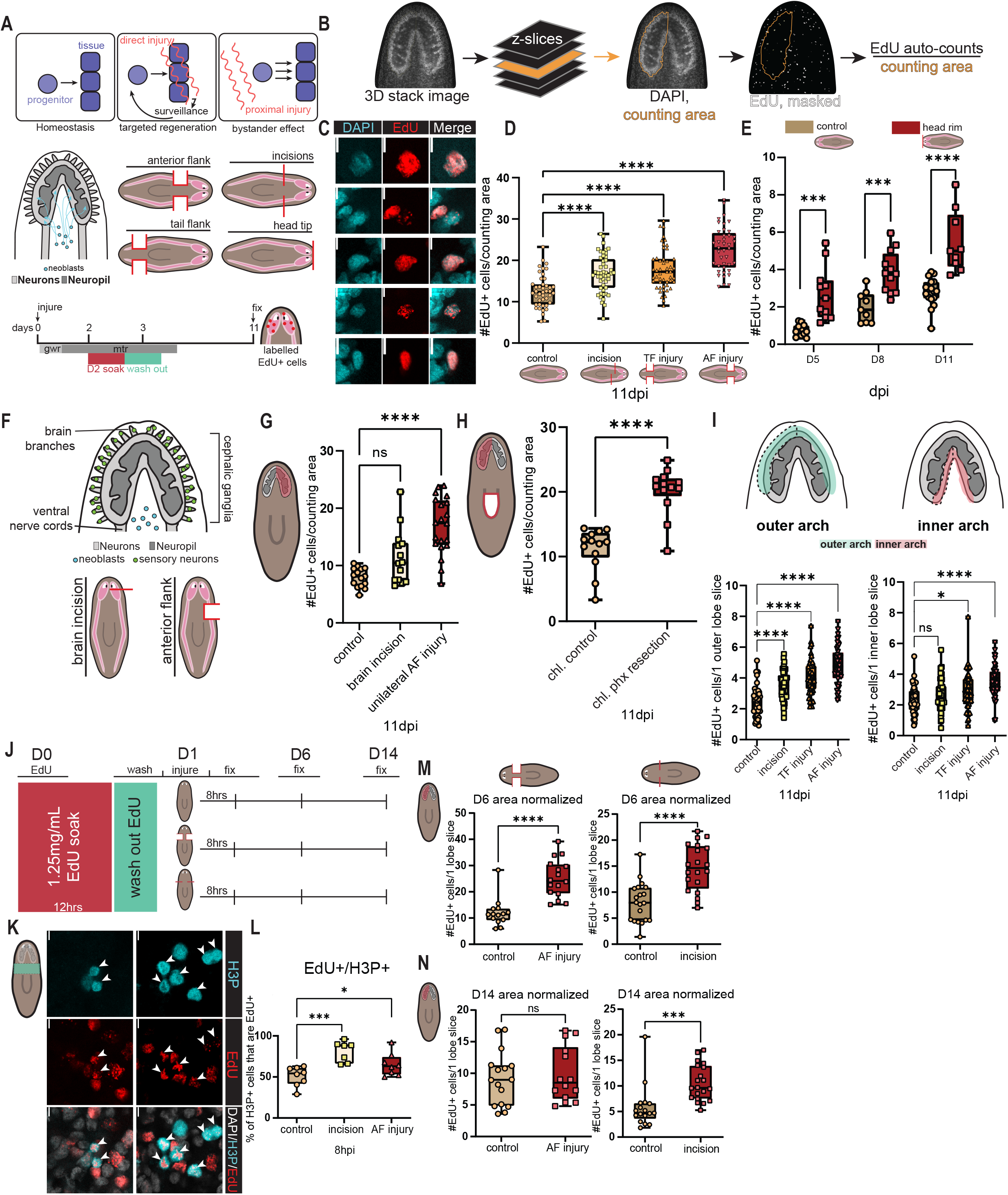
Injuries outside the brain increase incorporation of new neurons in the brain. (**A**) Cartoon of regeneration hypotheses, neoblast location, and integration. EdU delivery, injuries for D-F. (**B**) Semi-automated counting/brain-slice. (**C**) EdU+ brain cells; scale-bar, 5μm. (**D**) EdU+ cells/cephalic ganglion Z-slice (counting-area), one-way ANOVA, Dunnett’s Test correction, n≥45 (Table S1). (**E**) Timecourse: EdU+ cells per brain-slice per condition, unpaired Student’s t-test, n≥9. (**F**) Schematic and injuries for G. (**G**) EdU+ cells/cephalic ganglion Z-slice, one-way ANOVA, Dunnett’s Test correction, n≥14. (**H**) Pharynx removal, EdU+ cells/cephalic ganglion Z-slice, one-way ANOVA, Dunnett’s Test correction, n≥13. (**I**) Outer-versus-inner cephalic ganglion EdU+ nuclei/ Z-slice. Black dotted-line: counting-area. one-way ANOVA, Dunnett’s test correction, n≥44. (**J**) EdU schematic: H-K. (**K)** H3P+, EdU+ cells; scale-bar, 5μm. (**L**) Percentage of H3P+ cells that were EdU+, one-way ANOVA, Tukey-Kramer test correction, n≥6. (**M, N**) EdU+ cells at 6 dpi (**M**) 14 dpi (**N)** (unpaired Student’s t-test, n≥14). (**D-E, G-I, J-N**) Symbols: individual animals, scored blind; box plots with mean +/-SD; *p<0.05, **p<0.01, ***p<0.001, ****p<0.0001, n.s. = p>0.05.

The target-blind hypothesis suggests that large injury amplifies progenitors in a region-appropriate manner, but lacks specificity regarding the precise identity of lost tissues. A facet of this hypothesis is the “bystander effect,” where coincidence of injury-triggered neoblast proliferation with a progenitor specification location generically amplifies production of local progenitors, even for tissues that are not missing (LoCascio et al., 2017). This is predicted to occur because the distributions of some fate-specified neoblasts are broader than their target tissues. Here we find that planarian brain, specific CNS neurons, ventral nerve cords (VNCs), and the pharynx display this bystander effect. Our findings also indicate that progenitor responses to injury are stratified: the epidermis relies on recruitment of post-mitotic progenitors for blastema production; muscle shows recruitment of MTR-induced muscle progenitors to the wound; and peripheral neurons exhibit generic, increased regional cell addition without bias towards the wound itself. Overall, our findings indicate that the identity of missing tissues does not strongly dictate regenerative outcome, although the general process we describe could be coupled in principle with a surveillance process added for certain tissues. We suggest that a core process of baseline tissue turnover, positional information, and injury-stimulated proliferation serve as the primary determinants of cellular identity during wound repair.

## RESULTS

### Injuries outside of the brain increase incorporation of new neurons in the brain

We studied new-cell incorporation following injuries that are proximal to target tissues, but do not remove them. Neoblasts were labeled with F-ara-EdU (Neef and Luedtke, 2011) (thymidine analogue, hereby “EdU”) for 16 hrs, starting 48 hpi, capturing the MTR proliferative peak. No divisions occur directly within planarian mature tissues; EdU labels neoblasts, which produce EdU-labeled post-mitotic progenitors that can be tracked though migration and incorporation into mature tissues (Fig. 1A). Several injuries were utilized to assess for a brain bystander effect: anterior-flank, tail-flank, anterior-incision, and head-tip removal (Fig. 1A, S1A-B). The anterior-flank, anterior-incision, and head-tip injuries all cause proliferative responses in the animal anterior, with the incision leading only to a generic-wound-response proliferative peak and some residual elevated proliferation at 48 hpi (Wenemoser and Reddien, 2010). The tail-flank injury causes sustained proliferation (MTR-associated), but localized to the posterior.

EdU+ cells per cephalic ganglia z-slice were quantified with normalization to lobe area (Fig. 1B). Cephalic ganglia EdU+ cell incorporation significantly increased following anterior-flank, tail-flank, and anterior-incisions – injuries that did not directly remove cephalic ganglia cells (Fig. 1C-D). Anterior-flank injury drove the most substantial cell incorporation increase, whereas incision and tail-flank injuries showed similar, modest increases (Fig. 1D). This suggests that any proliferation increase occurring where brain progenitors are specified can lead to increased new-cell incorporation into the uninjured brain. Although these injuries could damage distal brain axons and in principle activate surveillance mechanisms, nerve cord injury did not cause overt acute brain cell death at 4 hpi (Fig. S1E-F). We assessed EdU+ cell incorporation into the brain following a head-tip injury (Fig. 1A), an injury occurring in an area devoid of neoblasts (Baguna, 2012; Newmark and Sánchez Alvarado, 2000) or VNCs, but capable of inducing MTR-proliferation at 48 hpi (Fig. S1C-D) (Wenemoser and Reddien, 2010). This injury also caused a significant increase in the number of EdU□ cells incorporated into the brain (Fig. 1E) – suggesting that the bystander effect in the brain was not the result of surveillance-based detection of damage to nerve cords or progenitors.

Bisecting the cephalic ganglia by incision severs many axons without removing neurons (Fig. 1F). If projection damage drove the observed responses, this injury should elicit a similar effect. Brain-incision, however, resulted in fewer prepharyngeal mitoses 48 hpi than did unilateral anterior-flank injury (Fig. S1G). Accordingly, brain-incision injury did not result in significantly elevated progenitor incorporation, whereas anterior-flank injury did (Fig. 1G). Pharynx resection causes little injury to brain axons/neurons; some cephalic ganglia axons innervate the pharynx, most pharynx neurons connect to VNCs (Cebrià, 2008; Okamoto et al., 2005). Pharynx resection substantially increases neoblast proliferation in the pre-pharyngeal region (Adler et al., 2014), where neuron progenitors are generated, and significantly increased EdU+ cell incorporation into cephalic ganglia (Fig. 1K). These data suggest that axonal or sensory neuron damage is unlikely to be a major contributor to the observed brain bystander effect. Instead, these data support a model in which injury-induced neoblast proliferation in progenitor specification zones increases cell production, even for cell types not lost to injury.

To ascertain whether the brain bystander effect was primarily driven by new neuron or glia incorporation, we analyzed the predominantly neural inner and outer arches of cephalic ganglia (Fig. 1A, I) (Roberts-Galbraith et al., 2016; Wang et al., 2016). Both arches displayed increased cell-incorporation in response to anterior-flank and tail-flank injuries (Fig. 1I). However, only the outer arch showed increased incorporation in response to anterior incisions, and overall showed greater incorporation responses to all injury types than did the inner arch (Fig. 1I). We conclude that the observed brain bystander effect involves increased incorporation of new neurons.

### The generic wound response contributes to a bystander effect

Anterior incisions do not robustly induce the MTR, and tail-flank injuries bias MTR-associated proliferation to the posterior. How then, did these injuries cause increased cell incorporation into uninjured brains? Fraction-of-mitoses-labeled experiments estimate a 6 hr neoblast G2 phase after feeding stimulation; generic-wound-response proliferation likely accelerates G2 rather than S-phase (Newmark and Sánchez Alvarado, 2000). Whereas the generic-wound response is initiated hours following injury, mitotic activity after incisions does not return to baseline by the time the MTR would normally occur (Wenemoser and Reddien, 2010). Because the generic-wound response elevates proliferation body-wide (Baguñà, 1976; Saló and Baguñà, 1984; Wenemoser and Reddien, 2010), EdU labeling at 48 hpi may have captured divisions contributing to increased brain incorporation following anterior incisions and tail-flank cuts.

To determine whether the generic-wound response can induce a bystander effect, we delivered EdU for 12 hours prior to injury (Fig. 1J). Roughly 50% of the H3P-labeled mitoses were EdU+ in controls, compared with ∼70% and ∼65% for anterior incision and anterior flanks (Fig. 1K-L), suggesting that pre-injury EdU-labeling captured cells contributing to the generic-wound response. Following anterior-flank injury of animals labeled in this manner, a significant increase in new-cell incorporation into a representative z-slice of the brain occurred by 6 dpi (Fig. 1M), with this effect absent by 14 dpi (Fig. 1N). In response to anterior-incision, a significant increase in EdU+ cell incorporation into the brain occurred at 6 dpi (Fig. 1M) and was sustained at 14 dpi (Fig. 1N). The longer perdurance of elevated EdU+ numbers is consistent with the observation that the generic-wound response does not induce a global and sustained increase in apoptosis, as does the MTR (Pellettieri et al., 2010). These data suggest that any proliferation occurring where neural progenitors are specified – whether the tissue is injured or not, or whether substantial tissue loss occurs or not – is sufficient to drive increased incorporation of new brain neurons.

### Increased progenitor and mature neuron formation in response to injuries outside of the brain

Some brain neuron classes also exist in the VNCs; in principle, injury to VNC neurons could trigger a surveillance process leading to elevated incorporation in the brain. Although this would involve increased new-cell incorporation into uninjured regions, we sought to investigate neural cell-types with restricted localization (Fig. 2A). dd_17258+ neurons are candidate sensory neurons localized peripherally in the head (Fincher et al., 2018). There was a significant increase in EdU+; dd_17258+ cells following anterior-flank injury only (Fig. 2B-C). This effect was observed when considering the absolute numbers of events, and when considering the number of EdU+; dd_17258+ cells as a percentage of the total dd_17258*+* population (Fig. S2A).

**Figure 2:**
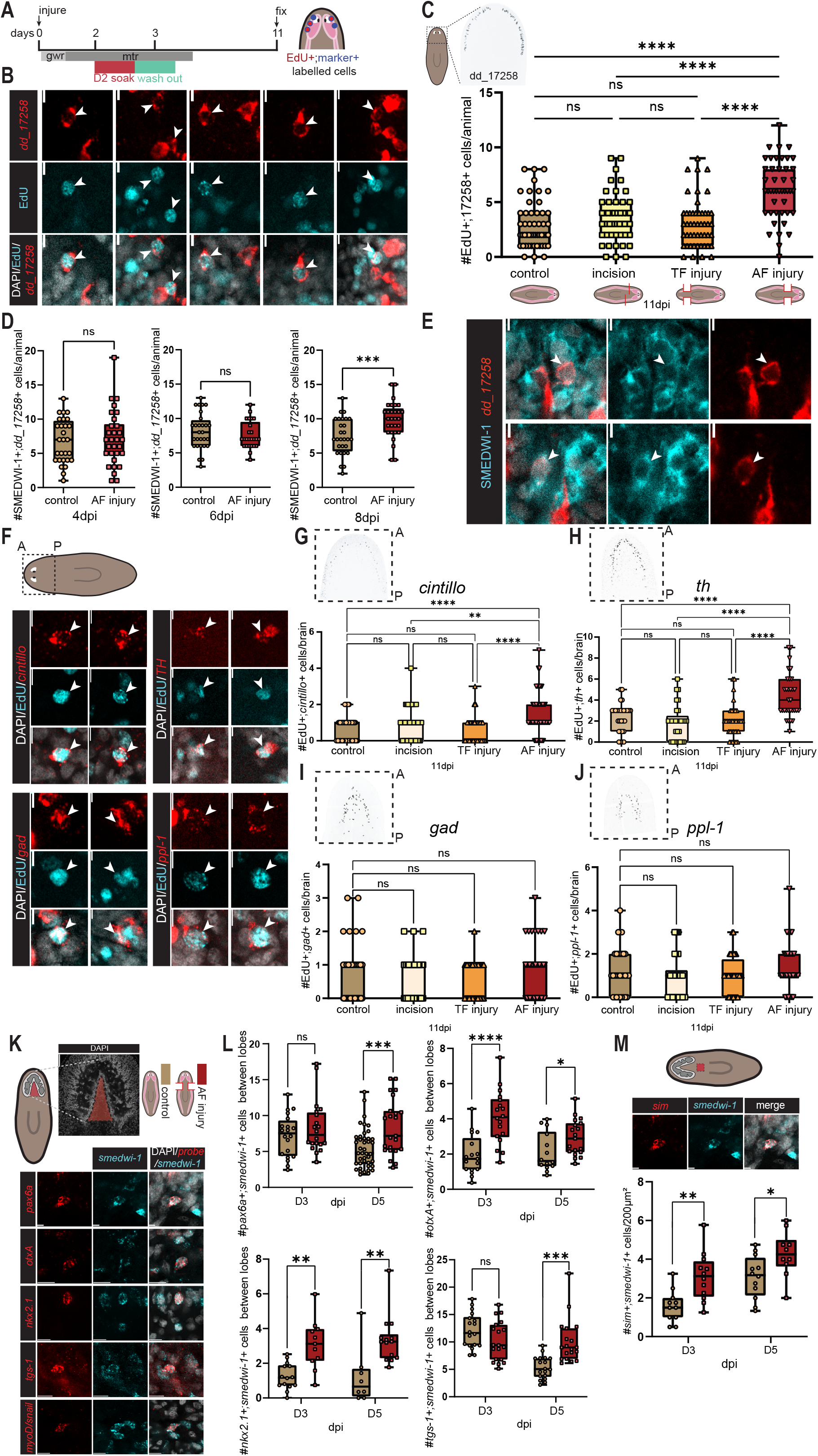
Increased incorporation of mature neurons in response to injuries outside the brain. (**A**) EdU delivery; injuries B, G-J. (**B**) dd_17258*+*; EdU+ cells (arrows), 5μm scale-bar. (**C**) EdU+; dd_17258+ cells/ animal (one-way ANOVA, Tukey-Kramer test correction, n≥45). (**D**) SMEDWI-1+; dd_17258*+* cells at 4, 6, and 8 dpi (unpaired Student’s t-test, n≥18). (**E**) SMEDWI-1+; dd_17258 + cells. 5μm scale-bar. (**F**) Cartoon: counting-area. A, anterior; P, posterior. *probe*+; EdU+ cells, 5μm scale-bar. (**G-J**) *probe+*; EdU+ cells (one-way ANOVA with Tukey-Kramer test correction, n≥29). (**K**) Counting-area (bar, 50μm); *probe+*; *smedwi-1+* cells; scale-bar, 10µm. (**L**) *probe+*; *smedwi-1+* cells at 3, 5 dpi in counting-area, n≥8. (**M**) Counting area. *sim+*; *smedwi-1+* cells at 3, 5 dpi in 200μm^2^ prepharyngeal region (unpaired Student’s t-test, n≥10). (**B, D, G-M**) Symbols: individual animals, scored blind; box plots with mean +/-SD; *p<0.05, **p<0.01, ***p<0.001, ****p<0.0001, n.s. = p>0.05.

To independently assess the bystander effect, we utilized SMEDWI-1 protein perdurance. *smedwi-1* transcription ceases after cell-cycle exit but SMEDWI-1 protein can be detected in neoblast descendants for several days, allowing detection of newly differentiated cells (Scimone et al., 2010; Wenemoser and Reddien, 2010). The increase in SMEDWI-1+; dd_17258+ neurons eight days following anterior-flank injury (Fig. 2D-E) was consistent with the EdU-labeling experiment, supporting a bystander effect.

We assessed four additional brain cell types, counting cell-incorporation events blind to condition. *cintillo*+ neurons (dd_11901), described as sensory (Oviedo et al., 2003), and *TH+* (*tyrosine hydroxylase*, dd_16581) dopaminergic neurons (Nishimura, 2007; Nishimura et al., 2011), both showed increases in cell incorporation following anterior-flank injury – similar to dd_17258*+* neurons (Fig. 2F-H). Total dd_17258*+* cell number showed only a modestly significant increase (Fig. S2B) and *TH+* neuron number did not change (Fig. S2C); however, cell numbers are also expected to be pushed downwards by an MTR-associated cell-death increase following this injury. *gad*+ cells (*glutamic acid decarboxylase*, dd_12653), inhibitory GABAergic neurons (Nishimura, 2008), and *ppl-1*+ (*pyrokinin prohormone like-1*, dd_14095) neurons (Collins et al., 2010) did not show increased cell-incorporation under the conditions described above (Fig. 2F, I-J), or after a medial-anterior hole-punch injury (Fig. S2D). *gad*+ and *ppl-1*+ cells reside in the medial brain, and this lack of response is consistent with the lower cell incorporation of the inner cephalic ganglia arch (Fig. 1I). Whether these cell types exhibit smaller or differently timed bystander effects, or differ in turnover dynamics, remains to be determined. Together, these results indicate that the bystander effect of the nervous system is a broad attribute of the planarian injury response.

To directly interrogate progenitors, we utilized RNA probes to neural (primarily transcription factor (TF)-encoding) gene transcripts in *smedwi-1+* cells: *pax6a* (dd_17726), expressed in a broad class of neural progenitors (Scimone et al., 2014), *otxA* (dd_14633), associated with cephalic ganglion neurons and photoreceptors (Lapan and Reddien, 2011; Scimone et al., 2014), *nkx2.1* (dd_13898), a TF expressed in brain neurons (Scimone et al., 2014), *sim* (dd_17731), required for the regeneration of several neuron types (Cowles et al., 2013; Scimone et al., 2014), and *tgs-1* (dd_10988), which marks neural-fate-enriched neoblasts (King et al., 2024; Raz et al., 2021; Zeng et al., 2018). None of these TFs are wound-induced, but instead they increase in transcript level during the MTR of head regeneration, when head progenitors are generated (Fig. S2E). We counted neoblasts between cephalic ganglia (Fig. 2K), except for *sim*+ cells, which were counted pre-pharyngeally as little *sim* expression was observed between cephalic ganglia (Fig. S2F). Because *smedwi-1* transcription ceases as neoblasts exit the cell cycle, *transcription-factor+; smedwi-1+* cells are newly specified neural progenitors. There was a significant increase in *transcription-factor+; smedwi-1+* and *tgs-1+*; *smedwi-1+* cells by 5 dpi following anterior-flank injury in all cases, with *sim+, nkx2.1+*, and *otxA+* cells also showing a significant increase at 3 dpi (Fig. 2K-L). This further suggests that direct injury to the brain is not necessary for amplification of several neural progenitors.

### The bystander effect is observed in the planarian nerve cords in response to proximal injury

To cause significant injury without impacting VNCs, we removed the pharynx. Pharynx resection causes a proliferative response (Adler et al., 2014), but this was proposed to be selective for pharynx progenitors through a surveillance program (Adler et al., 2014; Bohr et al., 2021). Prior work showed increased BrdU+ cell incorporation into VNCs following pharynx resection (LoCascio et al., 2017), but employed a dorsal incision for pharynx resection that could have contributed to the effect (Bohr et al., 2021). We excluded this possibility using two pharynx-resection methods (Fig. 3A): resection under 0.2% chloretone, and pharynx expulsion with 100µM sodium azide (Adler et al., 2014; Bohr et al., 2021; Shiroor et al., 2018).

**Figure 3:**
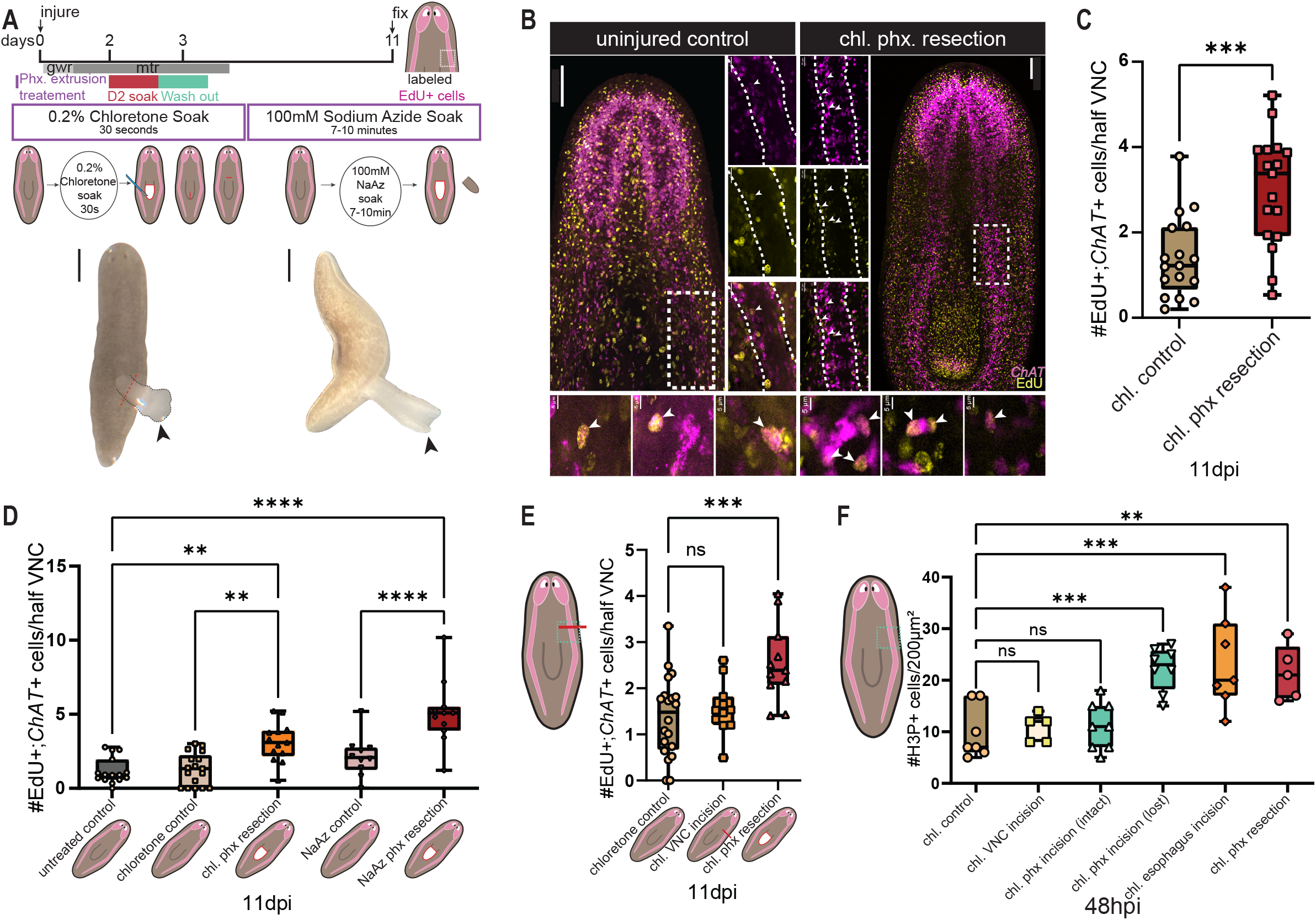
The bystander effect is observed in planarian nerve cords in response to proximal injury. (**A**) EdU delivery; injuries for B-E; 0.2% chloretone, 100µM sodium azide delivery, pharynx-resection. Black arrows: pharynx posterior. 300μm scale-bars. (**B**) Representative animals (100μm scale-bar). Dashed line: counting-area. Zoom: white-outlined VNC (20μm scale-bar); *ChAT*+; EdU+ cells (5μm scale-bar). (**C)** *ChAT+*; EdU+ VNC cells (unpaired Student’s t-test, n≥17). (**D)** 11 dpi *ChAT+;* EdU+ VNC cells (one-way ANOVA, Tukey-Kramer test correction, n≥10). (**E**) *ChAT+;* EdU+ VNC cells: direct-versus proximal-injury (one-way ANOVA, Dunnett’s test correction). (**F**) #H3P+ cells in 200uM^2^ area coinciding with quantified VNC in B. one-way ANOVA, Dunnett’s Test correction, n≥5. (**C-F**) Symbols: individual animals, scored blind; box plots with mean +/-SD; *p<0.05, **p<0.01, ***p<0.001, ****p<0.0001, n.s. = p>0.05. Chloretone: “chl.”, sodium-azide: “NaAz”.

In either treatment pharynx removal occurs without injury to epidermis, muscle, or parenchyma. Following pharynx removal, we labeled neoblasts with EdU at 48 hpi and assessed new-cell incorporation of new *ChAT*+ VNC neurons (Fig. 3B, S3). There was a significant increase in incorporation into the VNCs after chloretone-mediated resection (Fig. 3C-D), and even more strikingly in the sodium azide-treated pharynx-removed animals (Fig. 3D). This suggests that large injury outside of the VNCs causing increased proliferation can increase new VNC neuron incorporation, regardless of pharynx removal method (Fig. S3C).

To determine if axon damage could drive the observed VNC-bystander effect, we incised near the esophagus – the pharynx-intestine junction (Fig. S3A). Esophagus incision did not cause an increase in new-cell incorporation into VNCs, suggesting that axon damage did not significantly contribute to the observed effects (Fig. S3B). Furthermore, we directly incised the VNC itself, severing the nerve tract. VNC bisection did not result in a significant increase in new *ChAT*+ VNC neurons compared to controls, unlike pharynx resection (Fig. 3E). Notably, injury outside the VNCs that elevates proliferation increased VNC incorporation more robustly than did direct VNC injury that did not elicit sustained proliferation (Fig. 3F).

We attempted to injure but not remove the pharynx by vertical incision in chloretone-treated animals (Fig. S3A). This resulted in a significant increase in new-cell incorporation in the VNC (Fig. S3B), but the incision caused a substantial number of animals (∼50%) to expel their pharynx. Animals with and without pharynx-expulsion showed distinct mitotic responses (Fig. 3F), likely contributing to the increased new VNC cell incorporation (Fig. S3B). Prior work has reported that pharynx incision does not cause an upregulation of *FoxA*+ pharynx progenitors (Bohr et al., 2021).

### Pharynx-proximal injury causes a bystander effect in the pharynx

Is the bystander effect specific to the central nervous system or a broad general attribute? The pharynx has been implicated in both surveillance-based regeneration (Adler et al., 2014; Bohr et al., 2021) and as susceptible to a bystander effect (LoCascio et al., 2017). Injuries were designed to not impinge upon the pharynx and to induce: proliferation near the pharynx (pharynx-flank injury), smaller body-wide proliferation (trunk incisions), distant proliferation (head-wedge injury), or axon severing near the esophagus (esophagus-incision injury) (Fig. 4A, S4A-C). We assessed EdU-labelled cell incorporation across a single pharynx z-plane (Fig. 4A); only pharynx-flank injury, the injury causing increased proliferation near the pharynx, elevated pharyngeal EdU-labeled cell incorporation (Fig. 4B).

**Figure 4:**
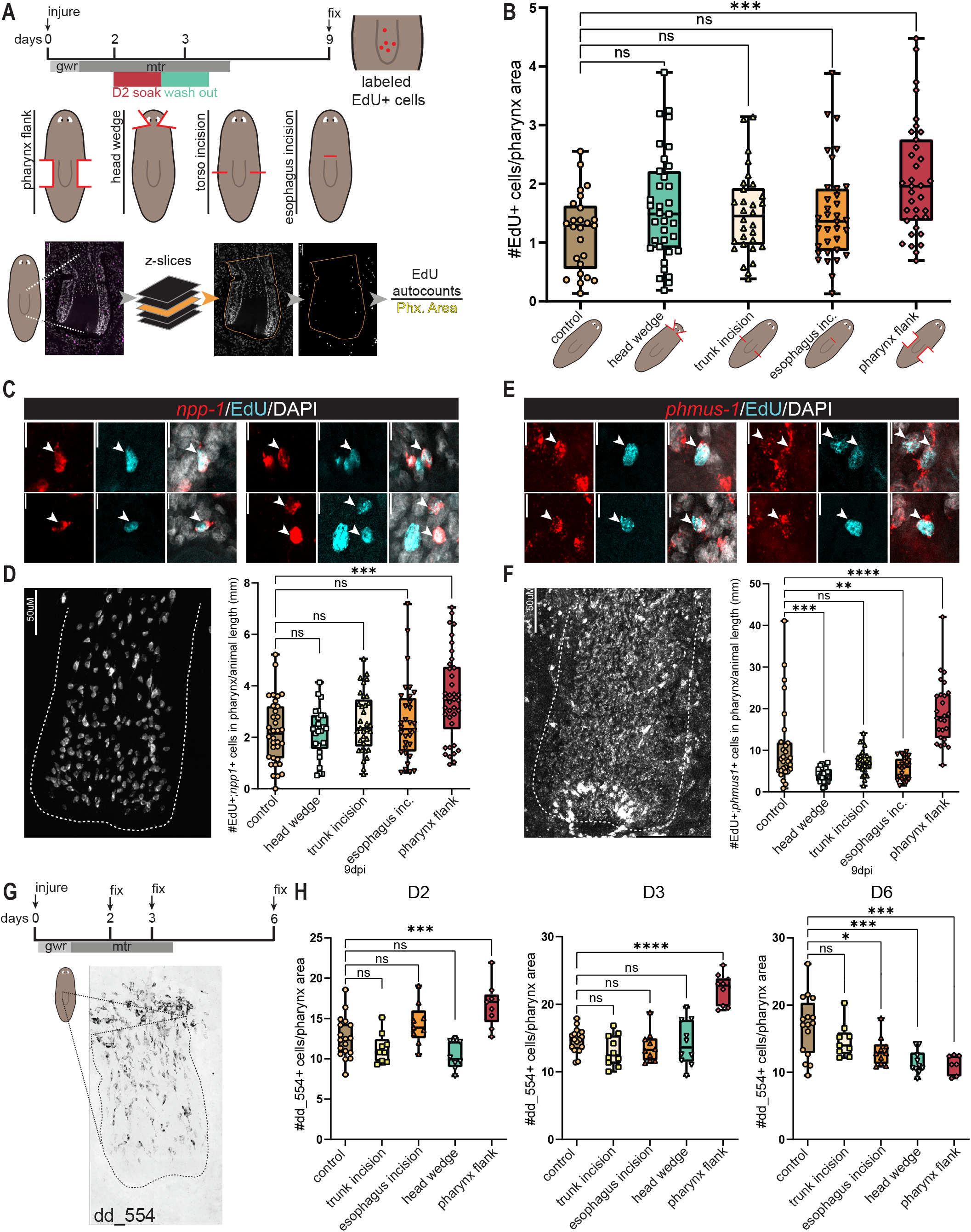
The pharynx responds to proximal injury with a bystander effect. (**A**) EdU delivery; injuries for B-F. Analysis for B. (**B**) EdU+ cells/ pharynx-slice (one-way ANOVA, Dunnett’s Test correction, n≥25). (**C**) *npp-1+*; EdU+ cells. 10μm scale-bar. (**D**) pharyngeal *npp-1* (dashed line). 50μm scale-bar. *npp-1+*; EdU+ cells/ fixed animal length (mm) for A (Fig. S4D) (one-way ANOVA, Dunnett’s Test correction, n≥34). (**E**) *phmus-1+*; EdU+ cells. 10μm scale-bar. (**F**) pharyngeal *phmus-1* (dashed line). 50μm scale-bar. *npp-1+*; EdU+ cells/ fixed animal length (mm) for injuries in A (Fig S4D). (one-way ANOVA, Dunnett’s Test correction, n≥21). (**G**) EdU delivery, injuries for H, dd_554*+* cells in pharynx (dashed line). (**H**) dd_554*+*; EdU+ cells/ pharynx area. (one-way ANOVA, Dunnett’s Test correction, n≥8). (**B, D, F, H**) Symbols: individual animals; box plots with mean +/-SD; *p<0.05, **p<0.01, ***p<0.001, ****p<0.0001, n.s. = p>0.05. (**D, F, H**) counted manually, scored blind.

There was a significant increase in EdU+; *npp-1+* (*neuropeptide precursor-1*, dd_2575) pharyngeal neurons and EdU+; *phmus-1+* (*pharynx muscle-1,* dd_8356) pharynx muscle only in response to pharynx-flank injury (Fig. 4C-F). Trunk incisions and head-wedge injuries modestly decreased *phmus-1+* incorporation, but this was not explored further (Fig. 4E-F). Because incisions fail to trigger sustained (MTR) proliferation, these data suggest that neither the muscle nor neural populations of the pharynx elevated incorporation because of damage to projections exiting/entering the pharynx. This further supports the model that pharynx-progenitor production can be upregulated following injury-induced proliferation in the medial region, regardless of pharynx-injury state. Increased cell incorporation led to a modest increase in total *npp-1+* cell numbers following pharynx-flank injury (Fig. S4D); likely reflecting a transient increase before cell-death compensation.

To interrogate pharynx progenitor responses, we assessed dd_554+ post-mitotic pharynx progenitors (Fincher et al., 2018; Zhu et al., 2015) (Fig. 4G). There was a significant increase in dd_554+ cells relative to pharynx size at two-and three-dpi for pharynx-flank injury; other injuries did not cause a significant change in numbers (Fig. 4H). At 6 dpi, the observed effect disappeared, consistent with the expected transiency of the increase in proliferation after injury.

Taken together, our data demonstrate that the pharynx is susceptible to the bystander effect: injuries that do not remove pharynx cells can upregulate new pharynx cell incorporation.

### New peripheral neuron incorporation is amplified outside the wound following large injury

Beyond testing for the bystander effect with injuries that do not directly impact regional tissues, we explored the specificity of cell-type production to wound location by studying more uniform tissues. Is new-cell incorporation into mature tissues for most-to-all planarian cell-types amplified in response to large injury? Epidermis-specialized neoblasts amplify less strongly at wounds than do neoblasts for other tissues (van Wolfswinkel et al., 2014), suggesting stratified progenitor responses. Unlike localized tissues such as brain neurons, VNCs, and pharynx, broadly distributed tissues such as epidermis, muscle, and peripheral neurons are formed in all blastemas. Does new cell production for broadly distributed tissues display different regenerative responses than for regional tissues? Furthermore, if production of broadly distributed cell types is amplified by injury are new cells specifically and exclusively targeted to the injured region? Decapitated animals were assessed for incorporation of 48 hpi-labeled EdU+ cells into peripheral neurons (*glipr-1*+, dd_210) (Fincher et al., 2018) and muscle (*colF-2*+, dd_702) within head blastemas (Fig. 5A). Tissues showed either a non-significant (*glipr-1*+) or a significant (*colF-2+*) increase of EdU+ cell incorporation into blastemas compared to intact heads (Fig. 5A). Decapitation, however, complicates interpretation of progenitor amplification dynamics because it causes rescaling of PCG-expression domains. We instead utilized an anterior-flank injury to limit anterior-posterior rescaling and analyzed a medial body-region defined by the pharynx width that was not infringed on by injury (Fig. 5B). Within this region we defined: “wound-proximal”, between the anterior-flanks, and “wound-distal”, immediately anterior to the flank injuries and extending to the head tip (Fig. 5B). “In-blastema” defines the wound location itself, restored by regeneration.

**Figure 5:**
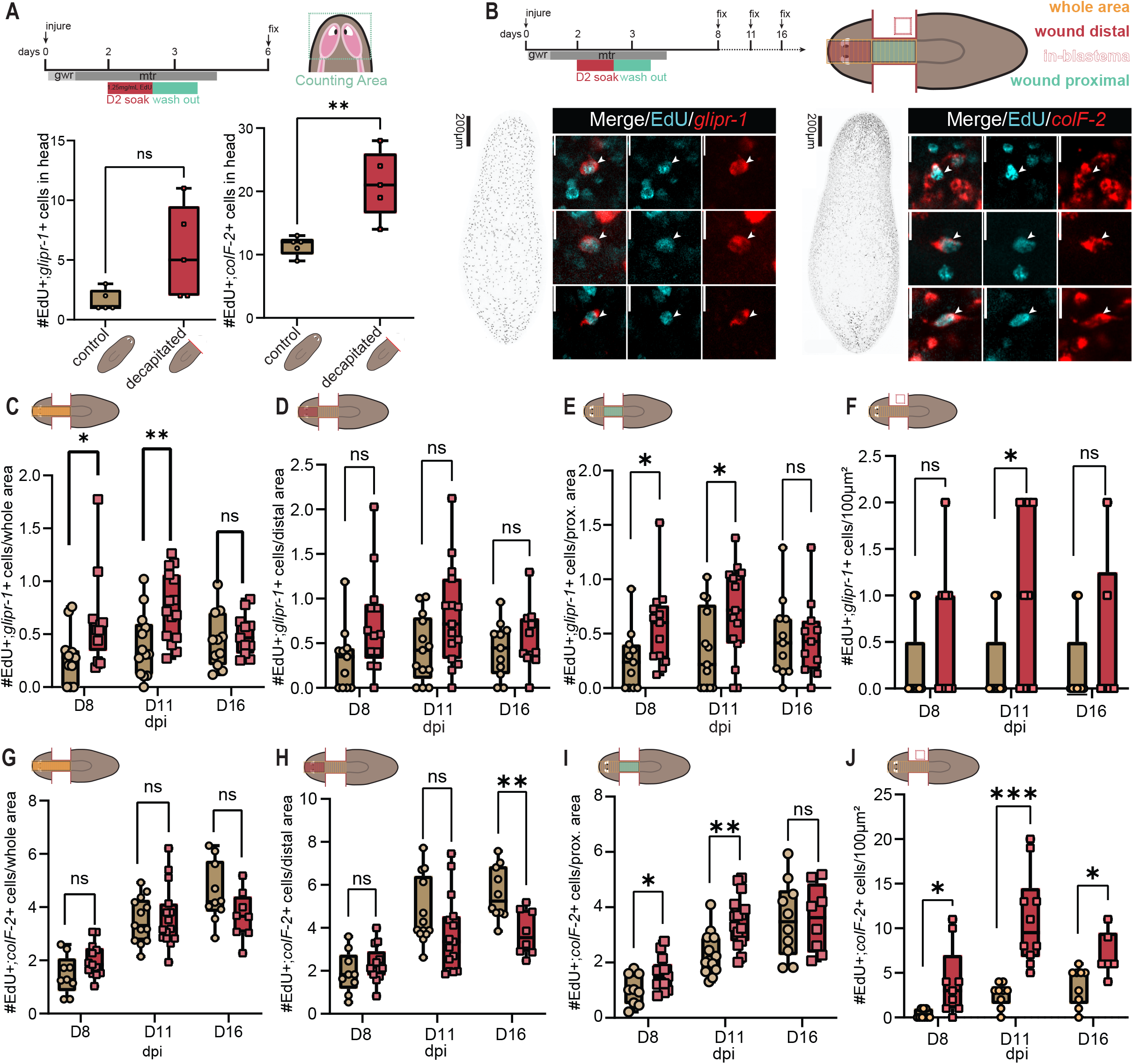
Broad amplification of peripheral neuron incorporation outside of injuries and local amplification of muscle incorporation following large injuries. (**A**) EdU delivery; injuries for A. EdU+; *glipr-1+* or EdU+; *colF-2+* cells/ counting-area (unpaired Student’s t-test, n≥5). (**B**) EdU delivery, injuries for B-J. Yellow: “whole-area” (wound-proximal and wound-distal), red: wound-distal, cyan: wound-proximal, dashed red box: in-blastema. EdU+; *glipr-1+* cells and EdU+; *colF-2+* cells. EdU+; *glipr-1+* cells in whole-area (**C**), distal-area (**D**), proximal-area (**E**), in-blastema (**F**) (unpaired Student’s t-test). EdU+; *colF-2+* cells in whole-area (**G**), in distal-area (**H**), in proximal-area (**I**), in-blastema (**J**) (unpaired Student’s t-test). (**C-F**) n≥11. (**G-J**) n≥8. (**A-B, C, D-F, H-J**) Symbols: individual animals, scored blind; box plots with mean +/-SD; *p<0.05, **p<0.01, ***p<0.001, ****p<0.0001, n.s. = p>0.05

At 8 and 11 dpi EdU+; *glipr-1*+ cell numbers (Fig. 5B-C) increased outside of the flank-injury region (including wound-proximal and-distal regions); by 16 dpi this difference was undetectable (Fig. 5C). No significant increase in incorporation of new *glipr-1+* neurons was observed in wound-distal areas (Fig. 5D), but was observed in wound-proximal (Fig. 5E). In a 100 μm^2^ blastema area there was a non-significant, and significant at 11 dpi, increase in the amount of new-cell incorporation into *glipr-1+* cells (Fig. 5F), although it was not substantially elevated compared to either wound-proximal or wound-distal areas. We suggest that injuries amplify incorporation of new *glipr-1+* cells within the blastema (where they are needed) as well as outside but proximal to the wound (where they were not removed or needed) to a similar degree. Whereas we cannot preclude a feedback-mechanism, considering injuries impact mature cells, these data point to a significant spatial imprecision in regenerative cell incorporation. Indeed, there was no observed difference in new-cell incorporation between the wound-proximal and-distal zones in injured animals (Fig. S5A).

### Incorporation of new muscle cells increases towards the wound, including in a wound-proximal region

Planarian muscle is widespread in the body and instructs regeneration outcomes by expressing positional information (Lander and Petersen, 2016; Scimone et al., 2016; Scimone et al., 2017; Witchley et al., 2013). Quantification of *colF-2+*; EdU+ cells (Fig. 5B) outside the wound showed no differences when comparing flank-injured and control animals (Fig. 5G, S5B). *myoD/snail+*; *smedwi-1+* muscle progenitors, showed detectable amplification only in the 3dpi blastema (Fig. S5C-F). In the wound-distal area there was no significant cell-incorporation difference between injured and control animals at 8 dpi, a non-significant decrease in incorporation following injury at 11 dpi, and a significant decrease in new-cell incorporation at 16 dpi (Fig. 5H). In the wound-proximal area there was a significant increase in new muscle incorporation at 8 and 11 dpi, which was no longer apparent 16 dpi (Fig. 5I). Analysis of a 100 μm^2^ blastema area showed a significant increase in incorporation of MTR-induced new *colF-2+* cells (Fig. 5J). These data suggest a biased incorporation of new muscle to sites within and proximal to the blastema, at the potential expense of progenitor incorporation (during turnover) into wound-distal sites. This represents a distinction in progenitor injury responses compared to peripheral neurons. At all timepoints in control conditions there was an anterior bias in new muscle incorporation, perhaps associated with the continual new-cell supply to the neoblast-devoid head-tip (Fig. S5B). By contrast, by day 11 post-flank injury this anterior bias was eliminated (Fig. S5B), suggesting reallocation of MTR-induced muscle progenitors from distal locations towards the wound. Nonetheless, like peripheral neurons, there was spatial imprecision in new muscle incorporation.

### Blastema epidermis is derived from pre-existing progenitors, with injury-associated amplification arriving after regeneration is complete

We suggest that amplification of epidermal neoblasts at large injuries, were it to occur, would contribute little to blastema formation. The planarian epidermis is derived from *zfp-1*-expressing (zeta) neoblasts, which produce post-mitotic progeny that mature during migration (Eisenhoffer et al., 2008; Tu et al., 2015; van Wolfswinkel et al., 2014; Wurtzel et al., 2017; Zhu and Pearson, 2016). Epidermal progenitor maturation requires ≥6 days, yet a functional epidermis-covered head regenerates within a week (Reddien et al., 2005; van Wolfswinkel et al., 2014). The epidermis is critical for the integrity the blastema, and actively participates in early blastema-outgrowth stages (Scimone et al., 2022). If six days are required for epidermal production, how can regeneration proceed so rapidly? An increase in post-mitotic epidermal progenitors has been observed at wounds (Wenemoser and Reddien, 2010), suggesting that pre-existing progenitors drive epidermis regeneration.

Dorsal and ventral epidermis were analyzed separately, because ventral epidermis cells have a significantly shorter half-life of 4.5 days (Lee et al., 2024). Following decapitation, neither epidermal plane showed significant – if any – incorporation of MTR-produced cells into mature epidermis of head blastemas (Fig. 6A-B). Indeed, 6 dpi, 4/5 animals showed no EdU+ cells in the ventral head epidermis at all, and 5/5 showed no EdU+ cells incorporation into the dorsal epidermis (Fig. 6A). By 11 dpi, ventral epidermis blastemas contain EdU+ cells (Fig. 6B, S6A-B). This further suggests that blastema epidermis formation does not rely on wound-induced progenitor proliferation.

**Figure 6:**
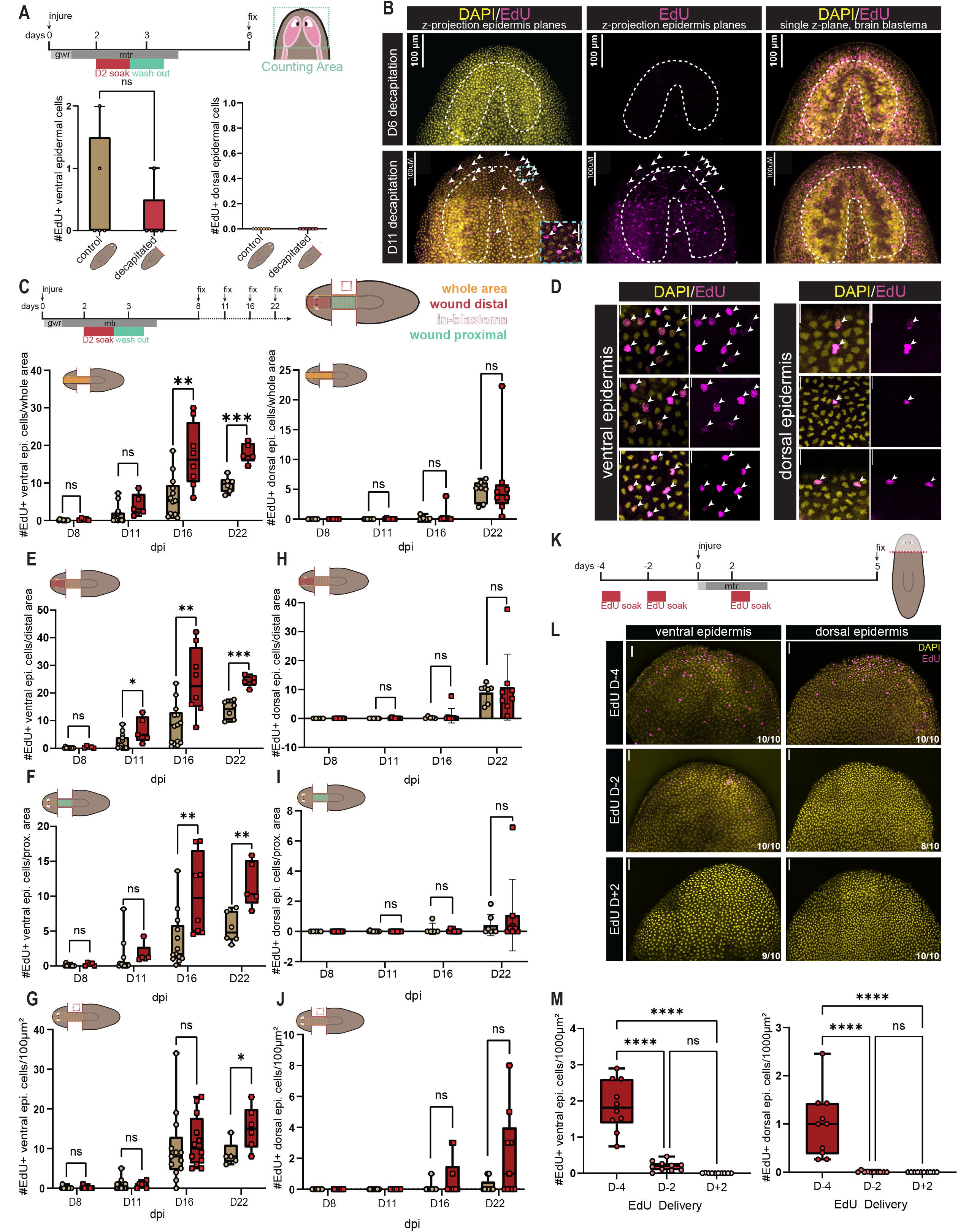
The blastema epidermis is derived from pre-existing post-mitotic progenitors, with injury-associated amplification arriving after regeneration is complete. (**A**) EdU delivery; injuries for A-B. EdU+ cells in dorsal or ventral epidermis/counting-area, 6 dpi. n≥5. (**B**) Ventral epidermis 6 and 11 dpi. Left/middle: Single z-plane ventral epidermis; right: single z-plane brain blastema (EdU-labeling control). Zoom: EdU+ epidermal cells (dashed blue box). Arrowheads: EdU+ epidermal cells, dashed line: brain. 100μm scale-bar. (**C**) EdU delivery, injuries for D-J; yellow: whole-area (wound-distal + wound-proximal), red: wound-distal, cyan: wound-proximal, dashed red box: in-blastema; EdU+ cells in whole-area (ventral and dorsal epidermis), unpaired Student’s t-test. (**D**) Ventral and dorsal epidermis. 10μm scale-bar. EdU+ ventral epidermis cells in distal-area (**E**), in proximal-area (**F**), in-blastema (**G**) (unpaired Student’s t-test, n≥5). EdU+ dorsal epidermis cells in distal-area (**H**), in proximal-area (**I**), in-blastema (**J**). (**K**) EdU delivery; injuries for L. (**L**) Ventral and dorsal epidermal blastema surfaces. Image corners: representative match. 200μm scale-bar (**M**) Quantification of L: EdU+ ventral and dorsal epidermal cells in blastemas, unpaired Student’s t-test, n≥10. (**C, E-F**) n≥5. (**H-J**) n≥6. (**D-J, M**) Symbols: individual animals, scored blind; box plots with mean +/-SD; *p<0.05, **p<0.01, ***p<0.001, ****p<0.0001, n.s. = p>0.05.

We observed essentially no incorporation into the ventral epidermis of either control or injured animals outside the injury-region until 16 dpi (Fig. 6C-D). Wound-proximal and wound-distal ventral epidermis behaved similarly: no detectable incorporation at 8 dpi, modest incorporation by 11 dpi, and an increase in EdU+ epidermal cells in the injured condition by 16 and 22 dpi (Fig. 6E-F). This suggests that injury amplified epidermal production, but only long after regeneration was complete and without wound-specific bias.

Additionally, whereas the anterior-flank injury caused an increase in new epidermal cells, from 16 dpi onwards there remained a robust bias towards anterior (wound-distal) incorporation (Fig. S6C). Incorporation into the blastema itself was not significantly elevated in injured animals compared to controls until 22 dpi (Fig. 6G). Together, these data suggest that there is eventual amplification of epidermal progenitor production in response to wounding, but no significant bias in the incorporation of newly produced cells to the wound. Considering the dorsal epidermis, there is a striking lack of response. The dorsal epidermis displayed almost no new epidermal cells until 16 dpi (Fig. 6C), demonstrated an anterior bias in incorporation (Fig. 6H-I, S6D), and did not appear to show significant bias towards incorporation in-wound (Fig. 6J), even by 16 or 22 dpi. There was an increased number of *agat-3*+ epidermal progenitors in all regions analyzed by 96 hpi (Fig. S6E-H), with the most substantial increase occurring in the blastema, where a modest increase was also observed at 24 hpi (Fig. S6H). At 0, 18, 24, 48, and 72hpi there was an anterior (wound-distal) bias in the number of *agat-3*+ cells, which was eliminated by 96 hpi (Fig. S6I), presumably a result of migratory recruitment.

We hypothesized that neoblasts that divided prior to wounding produced post-mitotic epidermal progenitors that are the source of new blastema epidermis. We delivered EdU in separate cohorts at 4-and 2-days prior to injury, as well as 2-days following injury, fixing animals 5 dpi (Fig. 6K). The ventral blastema epidermis at 5 dpi contained EdU+ cells for both EdU-delivery conditions prior to injury, but contained no EdU+ cells when delivery occurred during the MTR (Fig. 6L-M). The dorsal blastema epidermis only showed consistent EdU+ cell incorporation when EdU was delivered 4 days prior to injury.

These data, together with prior studies, suggest that the epidermis and its neoblasts respond to wounding in a distinct manner: *zfp-1*+ neoblasts do respond to injury by becoming amplified, at least to some degree, but unlike other tissues assessed, this amplification does not contribute to cell-type production during the time period of regeneration. Instead, the blastema epidermis relies on pre-existing post-mitotic progenitors, meeting the acute need for epidermal cells after injury (Fig. 7A).

**Figure 7:**
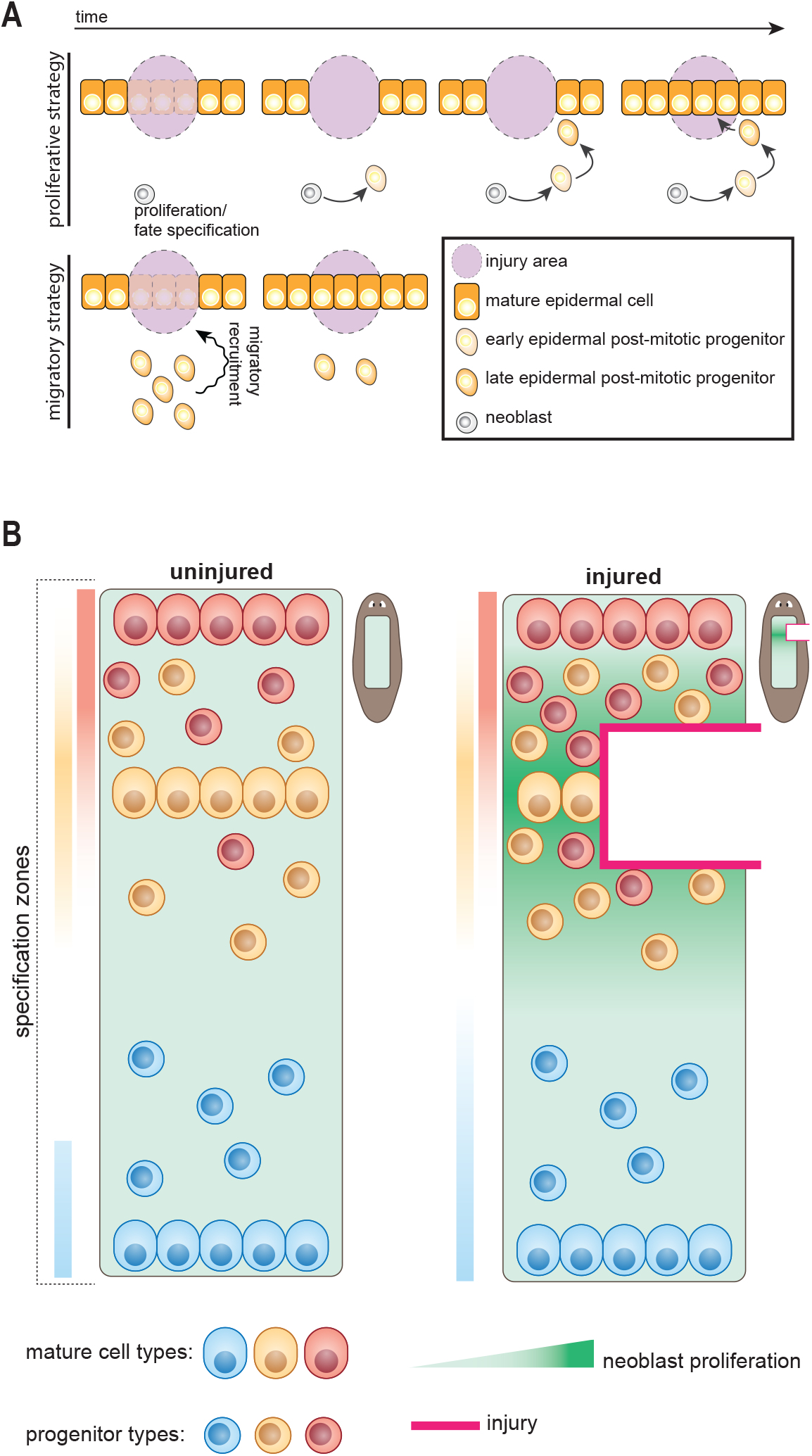
Model. (**A**) Neoblast proliferation driven regeneration (top), pre-existing post-mitotic progenitor driven regeneration (bottom). (**B**) Model: in uninjured conditions progenitors are continuously generated for tissue turnover in a spatially broad manner, based on positional information establishing progenitor specification zones (colored bars). Missing tissue wounds elicit sustained increases in proliferation (darker green). Coincidence of increased proliferation with progenitor specification zones results in amplified production of any cell type progenitors specified in that area.

## Discussion

How an animal replaces missing body parts and specific cell-types – regeneration specificity – is a poorly understood, central problem of regeneration. Planarians, with their highly plastic regenerative ability, provide a compelling context for study of this problem (Benham-Pyle et al., 2021; Fan et al., 2023; Owlarn and Bartscherer, 2016; Wurtzel et al., 2015). Our data support a model where any level of proliferation increase occurring where tissue-specific progenitors are specified is sufficient to increase incorporation of new cells into that tissue, regardless of the injury state of that mature tissue.

We suggest that planarians utilize a general mechanism to circumvent the complexity of sensing exactly which cell-types are missing (Figure 7B). Homeostatic cell turnover is continuous, and some progenitors are specified by constitutively active positional information in zones considerably broader than their target tissues. In this environment, local injuries can generically increase proliferation in neighboring stem-cells, yielding approximately the desired response: amplification of progenitor types suitable for the injured region. One consequence of this tripartite general turnover – positional information – injury-regulated proliferation mechanism is that it is noisy, resulting in the bystander effect. Because progenitor specification is broad, amplified progenitors will not constitute a perfect match to the identity of missing cell-types, and production of some cell-types that do not need to be regenerated at all is upregulated. An alternative solution for specifying production of missing cell types relies predominantly or exclusively on surveillance systems. Albeit more intuitive at first, such an approach to regeneration specificity would require a vast array of distinct surveillance mechanisms, each with suitable specificity/sensitivity to detect the exact type and degree of damage and dictating progenitor fate choices for 125+ adult cell types. We suggest that a turnover-based system combined with positional information and broad progenitor specification is a compelling solution that greatly simplifies the challenge of regeneration specificity.

In principle, some tissues might layer onto this generic mechanism a surveillance system that enhances the specificity and/or magnitude of the response provided. For instance, evolution could select for a mechanism that senses damage for tissues that are exposed to the environment and injured frequently (e.g., the pharynx). Such surveillance layered onto generic mechanisms is fully compatible with a bystander regenerative model. Much of our work utilized the number of new mature cells (EdU+ cells derived from neoblasts). This number could in principle be affected not just by progenitor-production increase, but also by other changes including increased progenitor survival above baseline. However, neoblasts and their progeny do not overtly contribute to homeostatic cell-death levels (Pellettieri et al., 2010); as would be expected if injury was causing a reduction in homeostatic progenitor death. Furthermore, neoblast proliferation increases correlated with observed bystander effects throughout this work, eye progenitors are amplified by injuries outside of the eye (LoCascio et al., 2017), and we observed increased numbers of many neural neoblast classes and pharynx progenitors in bystander experiments. These considerations support a model where increased regional neoblast proliferation increases new tissue production (Figure 7B).

The pharynx, implicated in both targeted and target-blind regeneration, is of particular importance for the problem of regeneration specificity. We challenged the pharynx for a bystander effect with pharynx-flank injuries to minimize positional information shifting. Decapitation has been used to study progenitor amplification behavior, but forces a trunk positional environment to transition into a head environment at the wound. It follows that is not conducive to the specification of pharyngeal progenitors, as the new PCG environment instructs neoblasts to make head cell-types. With medial injuries outside of the pharynx, we observed a pharyngeal-bystander effect – specifically for pharynx muscle, neurons, and post-mitotic dd_554+ pharynx-progenitors. Prior work showed pharynx removal results in the specific upregulation of *FoxA*+ pharynx progenitors (Bohr et al., 2021), demonstrating feedback-modulated targeted regeneration. Our data are compatible with this: the bystander effect observed in the VNCs after pharynx removal does not preclude the addition of signaling that enhances regeneration of the pharynx. In other words, pharynx removal could both induce an increase in mitotic activity that generically increases production of medial progenitors (for the pharynx and other, uninjured, cell-types) as well as trigger a signal that specifically boosts pharynx cell production. Our findings suggest, however, that injury to the mature pharynx is not strictly required to induce a pharynx-progenitor response.

We also considered how new-cell incorporation after injury might differ in uniformly-distributed tissues, such as muscle, peripheral neurons, and epidermis; any injury removing body regions would infringe on these cells. This could, in principle, generate conditions for different evolved progenitor behavior compared to regionalized tissues. Indeed, we found distinct responses to injury across these tissues. Where *glipr-1+* peripheral neurons demonstrated increased progenitor-based production in the entire anterior region with little to no bias towards the wound, incorporation of new *colF-2*+ muscle cells showed a wound-site bias. The candidate migratory recruitment process for muscle progenitors will be intriguing to investigate in the future. Near the wound, an increase in new-cell incorporation was observed for muscle, possibly associated with imperfect spatial amplification of this process towards the wound. Finally, epidermal regeneration relied exclusively on post-mitotic progenitors to form early blastema epidermis. MTR-induced epidermal progenitor amplification led to a wave of increased epidermal incorporation long after regeneration was completed and in uninjured regions. This points to a lack of specificity in the wound response, because epidermal progenitors amplified by wounding are not made to replace the missing epidermis in regeneration. Because epidermal progenitors require substantial production and maturation time, a pre-existing pipeline of near-mature precursors allows for rapid onset of acute repair – a benefit of turnover-mediated repair (Reddien, 2024) – preventing catastrophic delays in covering an outgrowth with skin (Fig. 7A). Without turnover, and therefore without a pre-existing pool of near-mature precursors, epidermal regeneration could not begin in less than six days, precluding rapid blastema formation. Notably, whereas post-mitotic epidermal progenitors have been shown to congregate at the wound (Wenemoser and Reddien, 2010) there is little evidence to determine whether these progenitors are responding to generic wound cues or to detection of epidermis damage itself. However, the late incorporation wave of MTR-induced epidermal progenitors was not wound-site specific.

The bystander effect could result in transient proportional changes between cells, but multiple important factors will impact relative cell numbers. The MTR triggers an increase in cell death that will push back on new cell addition outside of the wound. Furthermore, all tissues in planarians undergo turnover. It is unknown whether and how the incorporation rate of progenitors into a tissue impacts the death rate of that tissue. Regardless, once proliferation rates return to baseline after injury, turnover is expected to bring tissues, if out of proportion, back to steady state ratios (Reddien, 2024). The bystander effect itself could be a simple side effect of utilizing the proposed general solution to the challenge of regeneration specificity without relying on surveillance systems. However, it is also possible that the bystander effect yields desirable attributes. For instance, neighboring tissues suffering collateral damage from a wound could be replenished with new cells, helping ameliorate risk of transient dysfunction potentially caused by missing or mismatched cell populations. The bystander effect could also address transient progenitor depletion following migratory recruitment to wounds, or facilitate integration between wound-proximal and newly formed blastema tissues. Furthermore, several tissues exhibit wound-induced injury states of unclear function (Benham-Pyle et al., 2021). Investigation of how these states influence early progenitor behavior after proximal or direct injury, or other aspects of repair, will be of interest for future studies.

Based on the work presented here, contextualized within the field, we propose a general model for planarian regeneration specificity: the coincidence of constitutive positional information with elevated proliferation results in progenitor amplification from broad specification zones to ensure that wounds generally receive the progenitors they need (Fig. 7B). However, the specificity of this mechanism is not precise – a bystander effect is seen where increased production of cell-types occurs even for uninjured tissues. Even in the case of direct injury, incorporation is observed outside of the wound site. In axolotls, brain injury increased new neuron incorporation into a distal uninjured region (Amamoto et al., 2016) – raising the possibility that cell-type-production amplification outside of the wound, as described here, might occur broadly across regenerative contexts. We conclude that the identity of mature tissue itself has a limited impact on the specificity of progenitor responses to injury in planarian regeneration, with regeneration specificity instead being guided at least to a major degree by the broad factors of positional information and stem cell proliferation, factors extrinsic to the tissue itself.

## Materials and Methods

### Worm Care

*Schmidtea mediterranea* asexual clonal line CIW4 worms were maintained in static water culture systems containing Montjuïc salts (1.6□mmol/l NaCl, 1.0□mmol/l CaCl2, 1.0□mmol/l MgSO4, 0.1□mmol/l MgCl2, 0.1□mmol/l KCl and 1.2□mmol/l NaHCO3 prepared in Milli-Q water) (Cebrià and Newmark, 2005), “planarian water”. Animals were fed regularly with homogenized calf liver on a weekly or bi-weekly (every 2 weeks) basis and kept at 20°C in the dark outside of feedings. Prior to experimentation, animals were starved for 7 days. Animals used for experiments were between 3-5mm in length, size-matched within experiments, healthy, of a wild-type genotype, and not used in any prior experiment.

### Surgical Procedures

All surgical procedures and injuries were performed on planarian water-soaked filter paper on a cold Peltier block to immobilize the worms. Injures were performed guided by anatomical landmarks to ensure there was no impact on the tissue under analysis for a bystander effect.

### Chemical Treatments

Sodium azide pharynx removal was performed as previous described in the literature (Adler et al., 2014; Bohr et al., 2021; Shiroor et al., 2018). Following a 4-7min soak in 100µM sodium azide, animals were removed to a recovery dish with planarian water where those that retained their pharynx throughout the assay were maintained as a control group whereas those that lost it were considered for the injured group. Sodium azide has been known to suppress transcription and translation (Buchan et al., 2011), whereas in planarians, exposure suppresses mitoses in planarians for 24hrs, after which cells resume normal proliferation (Adler et al., 2014). Careful adherence to described protocols (Shiroor et al., 2018) can help mitigate the effect. Animals that retained their pharynges under sodium azide treatment were retained as controls to account for these potential effects. For the chloretone treatment, the animals were soaked for 30 seconds in a 0.2% w/v chloretone solution in planarian water, before being removed to wet filter paper on a cold Peltier block for pharynx resection. Pharynges either extruded on their own, and were then cut ∼3/4 of the way up from the tip towards the esophagus, or the animals were placed ventral side up and given a gentle massage to encourage the pharynx to slip out of the mouth, when it was resected as described. Chloretone control animals were soaked for 30 seconds before being placed directly into planarian water for recovery. Chloretone has known interactions with voltage-gated sodium channel type-II (Na_V_1.2), causing a dose-dependent and reversible inhibition (Kracke and Landrum, 2011). In mammalian cell culture, exposure to chloretone was shown not to alter Ca^2+^ dynamics of afferent discharging neurons or the shape of action potentials (Fischer, 2000). Nonetheless, control animals were soaked in chloretone for 30 seconds and allowed to keep their pharynges uninjured for direct comparison to chloretone injured animals.

### F-ara-EdU Labeling

F-ara-EdU (Vector Laboratories, CCT-1403) labeling was performed via soaking. Unless otherwise notes, all soaks were 16 hours in length in a solution of 1.25mg/mL F-ara-EdU prepared in planarian water. The animals were kept at 20°C in the dark for the duration of the soak. At the conclusion of 16 hours, the worms were washed 3x with a 1:1 solution of planarian water:5g/L Instant Ocean, and kept in 5g/L Instant Ocean until fixation with water changes every 3-4 days.

### Fixations and Whole-mount fluorescent in situ hybridizations

RNA probes were synthesized and whole-mount FISH was performed as previously described (King and Newmark, 2013; Pearson et al., 2009; Scimone et al., 2016). In brief, animals were killed in 5% NAC and fixed with 4% formaldehyde (w/v). Following deionized formamide bleaching, animals were treated with proteinase K (2ug/mL) and followed by a 4% formaldehyde incubation for 10 min each. Following overnight hybridization at 56°C with a 1:800 dilution of RNA probes in hybridization buffer (50% formamide, 5x SSC, 1 mg/ml yeast RNA, 1% Tween-20 and 5% dextran sulfate), samples were washed twice with, in order: pre-hybridization buffer, 1:1 pre-hybridization buffer:2xSSC, 2xSSC, 0.2xSSC, PBS with Triton-X (PBSTx). Blocking was performed with 5% Horse Serum and 5% Roche Blocking Reagent (Roche, 11921673001) PBSTx for DIG antibody, and with 5% Casein and 5% Roche Blocking Reagent for FITC antibody. In the event Roche Blocking Reagent was unavailable, blocking for both DIG and FITC antibody was performed with 5% Casein, 5% Horse Serum with no measurable difference in signal. All blocking solutions were filtered with a 0.22µM PES vacuum filter. Following overnight antibody incubation at 4°C, 2 hours of PBSTx washes were performed prior to tyramide development. For tyramide development animals were incubated for 10 minutes in borate buffer (0.1M boric acid, 2M NaCl, pH 8.5), followed by 10 minutes in borate buffer containing either fluorescein (1:1500), rhodamine (1:1000), or cyanine5 (1:300). Where necessary, 1.5 hours of peroxidase inactivation with 1% sodium azide was performed at room temperature, followed by 1.5 hours of washes prior to subsequent antibody incubations and tyramide developments.

SMEDWI-1 antibody (Guo et al., 2006) labeling was performed using a 1:1000 dilution in 10% Roche Blocking Reagent, and detected using the same tyramide detection as described above. H3P (Sigma-Aldrich, Anti-phospho Histone H3 S10 Antibody, clone 63-1C-8, rabbit monoclonal) antibody labelling was performed using a 1:300 dilution of the primary antibody in 5% Horse Serum, and detected using the same tyramide detection as described above.

For F-ara-EdU detection, whole-mount FISH was performed as described above until 4% formaldehyde was washed out. After proteinase K treatment but prior to hybridization, a CLICK chemistry reaction was performed by incubation in a solution of: 78.9µL PBS, 1.0µL 100mMCuSO_4_, 0.1µL 10mM Azide-Fluor 545 (MilliporeSigma 760757, resuspended in DMS), 20µL 50mM ascorbic acid (prepared fresh in water), for a total of 100µL — scaled up as needed. Samples were incubated in the dark on a shaker for 30 minutes, followed by 6 washes in PBSTx for a total of 45 minutes. Subsequently, FISH proceeded with hybridization as previously described.

### Microscopy and image analysis

Live images were acquired using a Zeiss Discovery V8 stereomicroscope with an AxioCam HRc camera. Fluorescent images were acquired on either a Leica Stellaris 5 WLL Confocal Microscope or a Leica SP8 Confocal Microscope, no comparisons were performed between images acquired on different microscopes. For single-channel quantifications of signal (i.e., H3P labeling), images were acquired at 20x magnification. For double-positive counting images were acquired at 40x or 63x magnification. Image analysis was performed using ImageJ/FIJI. All counting and co-occurrence of signal analyses were performed manually unless otherwise noted. Single-channel 3-dimensional computer assisted counting was performed using the Laplacian of Gaussian detector built into the TrackMate plugin in (Ershov et al., 2022), single-slice single-channel counting was performed using FIJI’s built-in Analyze Particles feature. In single-slice counting, the slice was selected based on >3 anatomical landmarks to approximate the same location between animals. For the brain, this consisted of selecting a slice that fulfilled the following criteria: (1) just above the ventral nerve cord connection to the brain, in the slice immediately above top-most layer of VNC visibility, (2) the connection between the two lobes must be open and free of nuclei (ie., neither too dorsal nor too ventral), and (3) the neuropil of the selected lobe must be clearly visible, and nuclei dense rim regions clearly defined. For single-slice counting in the pharynx the slice was selected that matched the following criteria (1) roughly in the middle of the pharynx, (2) the esophageal opening was readily visible in the anterior, (3) the top of the pharynx cavity was visible on either side, (3) the posterior opening of the pharynx was visible and clearly defined, (4) there was continuity of the inner pharyngeal tube with sparse cells from the anterior to the posterior. For normalization over animal length, fixed worms that were mounted on coverslips for confocal imaging were imaged under a dissection microscope along with a ruler. ImageJ/FIJI was used for quantification of worm size, using known measurement provided by ruler. For normalization over pharynx size, area measured at widest and longest point of the pharynx. All co-occurrence of signal analyses were performed blinded to image condition and providence, file names were replaced with random number identifiers. For epidermal counting experiments where dorsal and ventral epidermis were differentiated, the cells of the dorsal-ventral boundary (DVB) were excluded based on morphology. For visualization, but not analysis, of the curved surface of the epidermis, images were processed using the ImageJ/FIJI plugin VolumePeeler (Gatica et al., 2023). Gemini (Google) was used exclusively as a coding assistant for troubleshooting and debugging scripts. All final code was manually reviewed, tested, and validated by the authors. N numbers for all conditions quantified for each figure panel are presented in Table S1.

### Quantification and Statistical Analyses

Statistical analyses were performed using Prism software (GraphPad Inc., La Jolla, CA). Differences between the means of two populations were evaluated using a Student’s t-test. For comparisons between the means of three or more populations, a one-way ANOVA with multiple comparisons was performed; where means were compared between all groups Tukey-Kramer test was performed, for comparisons between the means only to the control condition Dunnett’s test was performed. A summary table listing key details for each injury paradigm and regenerative outcomes is provided in Table S2.

## Acknowledgements

The authors would like to thank Thomas Cooke, Lucila Scimone, and Patrick Aoude for helpful discussions; Patrick Aoude for computational help; Caitlin Rausch for assistance with model graphic design; Troy Whitfield for statistical modeling advice; the W.M. Keck Microscopy Innovation Center Core facility for microscopy advice; Thomas Cooke, Conor McMann, Lucila Scimone, and Deniz Atabay for comments on the manuscript. The authors acknowledge financial support from NIH R35GM145345.

