## Supplemental Figures for "Lack of specificity of planarian progenitor responses to injury in regeneration"

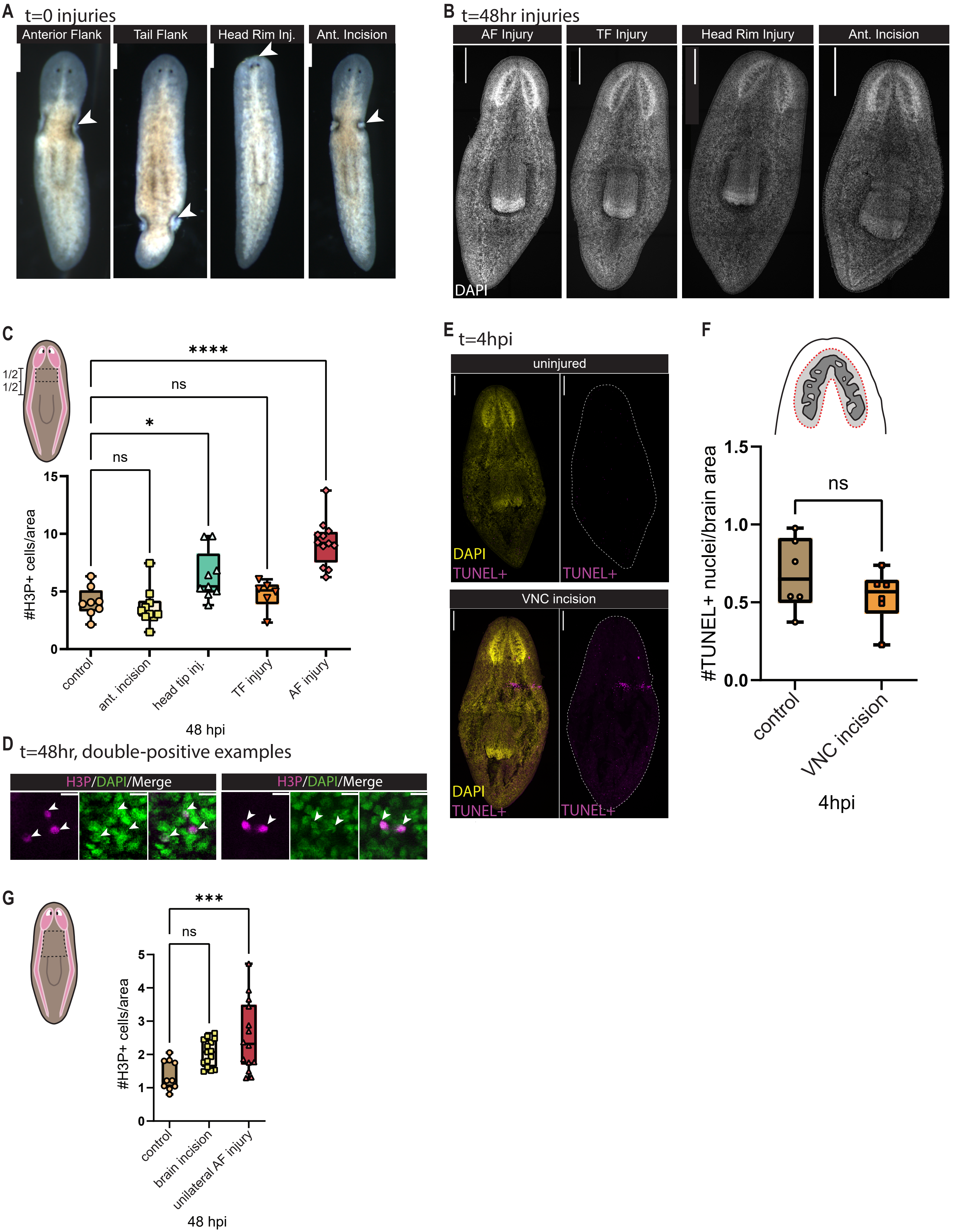
A t=0 injuries
Anterior Flank
Tail Flank
Head Rim Inj.
Ant. Incision
B t=48hr injuries
AF Injury
TF Injury
Head Rim Injury
Ant. Incision
DAPI
C
1/2
1/2
****
ns
*
ns
15
10
5
0
#H3P+ cells/area
control
ant. incision
head tip inj.
TF injury
AF injury
48 hpi
D t=48hr, double-positive examples
H3P/DAPI/Merge
H3P/DAPI/Merge
E t=4hpi
uninjured
DAPI
TUNEL+
TUNEL+
VNC incision
DAPI
TUNEL+
TUNEL+
F
1.5
1.0
0.5
0.0
#TUNEL+ nuclei/brain area
ns
control
VNC incision
4hpi
G
***
ns
5
4
3
2
1
0
#H3P+ cells/area
control
brain incision
unilateral AF injury
48 hpi

**Figure S1**

(**A**) Live images of injured animals immediately after injury; white arrows point to injury location. 200µm scale bar. (**B**) 20x confocal DAPI images representative of each injury, 48hpi. 300µm scale bar. (**C**) Number of H3P+ cells found in the prepharyngeal region as represented by the black dotted box in each injury condition. Counted with computer-aided, semi-automated methods, counting area defined by ventral nerve cords laterally and as half the distance from base of brain to top of pharynx vertically (one-way ANOVA with Dunnett's Test correction, $n \geq 6$). (**D**) Representative examples of H3P+ DAPI cells counted as H3P+, 20x, 5µm scale bar. (**E**) Representative examples of TUNEL+ staining in control and injured animals, 20x, 200µm scale bar. (**F**) Number of TUNEL+ nuclei in brain as represented in E (unpaired Student's t-test, $n \geq 6$). (**G**) Number of H3P+ cells found in the prepharyngeal region as represented by the black dotted box. Counted manually, counting area defined by ventral nerve cords laterally, bottom of brain anteriorly, and top of pharynx posteriorly (one-way ANOVA with Dunnett's test correction, $n \geq 10$). (**C, F, G**) Symbols: individual animals, scored blind; box plots with mean +/- SD; *$p<0.05$, **$p<0.01$, ***$p<0.001$, ****$p<0.0001$, n.s. = $p>0.05$. Complete n-numbers in Table S1.

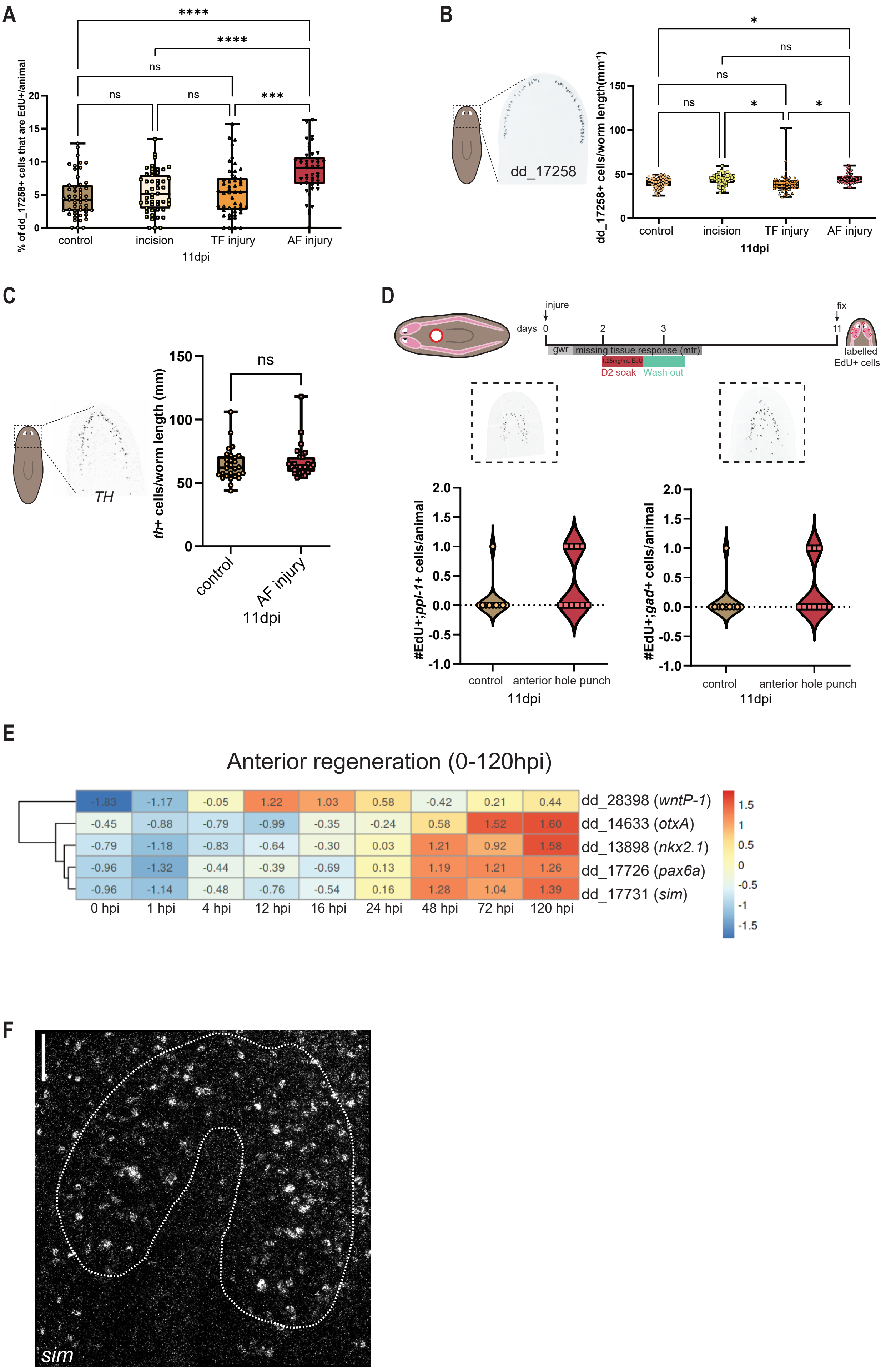

A
% of dd_17258+ cells that are EdU+/animal
****
****
ns
ns
ns
***
0
5
10
15
20
control
incision
TF injury
AF injury
11dpi
B
dd_17258
dd_17258+ cells/worm length(mm-1)
*
ns
ns
ns
*
*
0
50
100
150
control
incision
TF injury
AF injury
11dpi
C
TH
th+ cells/worm length (mm)
ns
0
50
100
150
control
AF injury
11dpi
D
days
injure
fix
0
2
3
11
gwr
missing tissue response (mtr)
D2 soak
Wash out
labelled EdU+ cells
#EdU+;ppl-1+ cells/animal
#EdU+;gad+ cells/animal
-1.0
-0.5
0.0
0.5
1.0
1.5
2.0
control
anterior hole punch
11dpi
control
anterior hole punch
11dpi
E
Anterior regeneration (0-120hpi)
-1.83 -1.17 -0.05 1.22 1.03 0.58 -0.42 0.21 0.44 dd_28398 (wntP-1)
-0.45 -0.88 -0.79 -0.99 -0.35 -0.24 0.58 1.52 1.60 dd_14633 (otxA)
-0.79 -1.18 -0.83 -0.64 -0.30 0.03 1.21 0.92 1.58 dd_13898 (nkx2.1)
-0.96 -1.32 -0.44 -0.39 -0.69 0.13 1.19 1.21 1.26 dd_17726 (pax6a)
-0.96 -1.14 -0.48 -0.76 -0.54 0.16 1.28 1.04 1.39 dd_17731 (sim)
0 hpi
1 hpi
4 hpi
12 hpi
16 hpi
24 hpi
48 hpi
72 hpi
120 hpi
1.5
1
0.5
0
-0.5
-1
-1.5
F
sim

**Figure S2**

**(A)** Computer-aided, semi-automated counting of total dd_17258+ cells per animal in each injury condition, a modest increase compared to control is seen only in the AF injury condition. Statistical significance assessed by one-way ANOVA with Tukey-Kramer test correction, n≥45.. **(B)** Total EdU+; dd_17258+ cells as a percentage of total dd_17258+ cells in each animal (one-way ANOVA with Tukey-Kramer test correction, n≥45). (**C**) Total th+ cells per animal in control and AF injury conditions unpaired Student's t-test, n≥10). (**D**) Schematic: EdU delivery relative to injury and fixation for, representation of hole-punch injury, probe+; EdU+ cells (unpaired Student's t-test, n≥28). (**E**) Heatmap showing activation of wound-induced gene expression 0-120 hpi in anterior-facing wounds of prepharyngeal region fragments. Only *wntP-2*, known to be wound-induced, shows increased activation before 48 hpi – all neural progenitor markers utilized in this study do not; data and visualization from (Wurtzel et al., 2015). (**F**) Z-projected 63x confocal image of *sim* expression in brain, cephalic ganglia outlined with dotted white line. (**A-D**) Symbols represent individual animals, data represented as box plot with mean +/- SD, *p<0.05, **p<0.01, ***p<0.001, ****p<0.0001, n.s. = p>0.05. Complete n-numbers in Table S1.

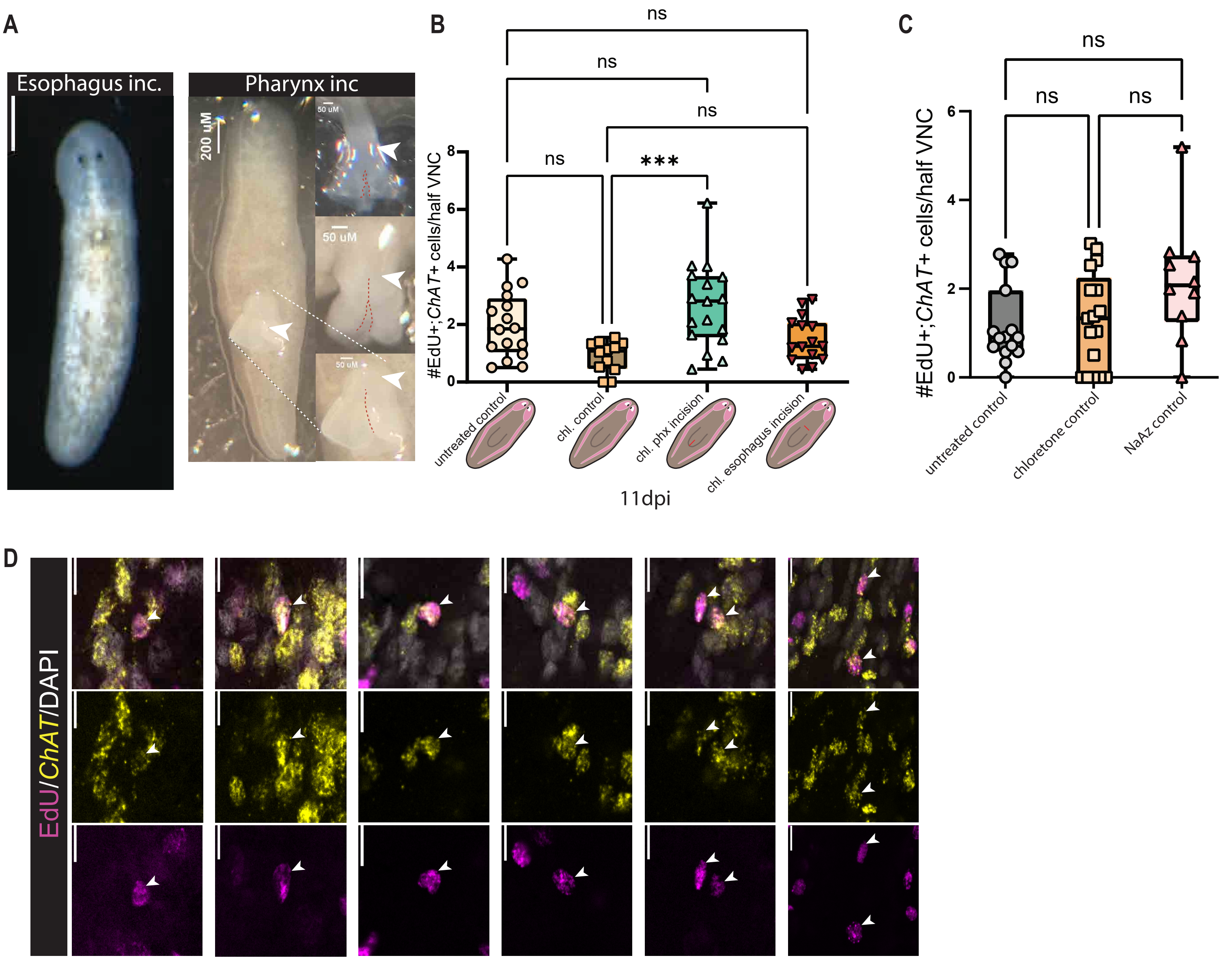

A
Esophagus inc.
Pharynx inc
200 uM
50 uM
50 uM
50 uM
B
ns
ns
ns
ns
***
#EdU+;ChAT+ cells/half VNC
0
2
4
6
8
untreated control
chl. control
chl. phx incision
chl. esophagus incision
11dpi
C
ns
ns
ns
#EdU+;ChAT+ cells/half VNC
0
2
4
6
untreated control
chloretone control
NaAz control
D
EdU/ChAT/DAPI

**Figure S3**

(**A**) Live images of injured animals immediately after injury, white arrows point to injury location. Zoom in view of pharynx in pharynx incision condition, with two additional examples. 200μm scale bar. (**B**) Number of EdU+; *ChAT*+ cells in each injury condition (one-way ANOVA with Tukey-Kramer test correction, n≥12). (**C**) Number of EdU+; *ChAT*+ cells in each chemical treatment condition, n≥10 (**D**) Representative images of *ChAT*+; F-ara-EdU+ cells counted in Figure 3. Each field of view grouped vertically. (**B-C**) Symbols represent individual animals, data represented as box plot with mean +/- SD, *p<0.05, **p<0.01, ***p<0.001, ****p<0.0001, n.s. = p>0.05. Complete n-numbers in Table S1.

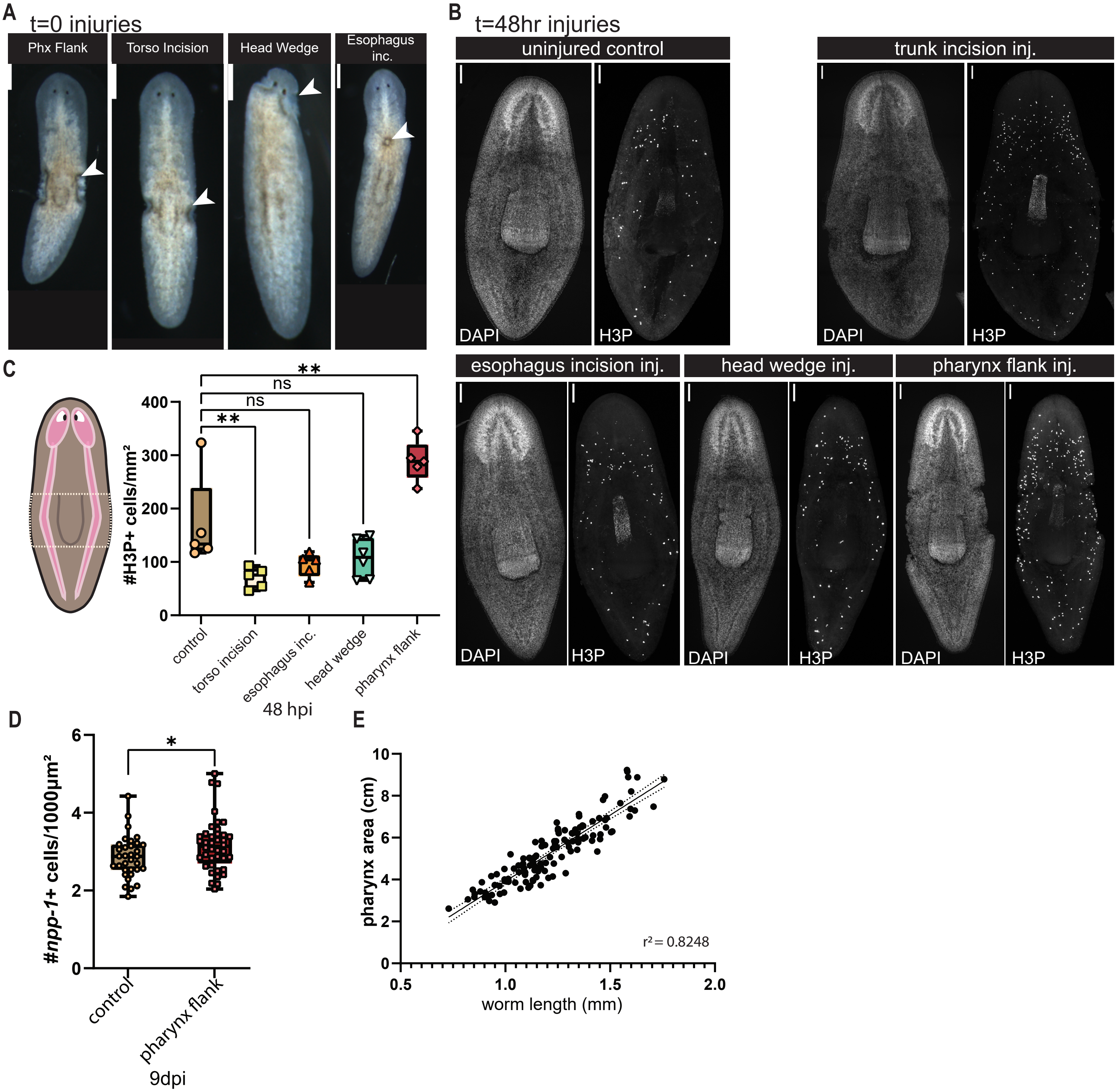
A
t=0 injuries
Phx Flank
Torso Incision
Head Wedge
Esophagus inc.
B
t=48hr injuries
uninjured control
trunk incision inj.
esophagus incision inj.
head wedge inj.
pharynx flank inj.
DAPI
H3P
C
#H3P+ cells/mm²
**
ns
ns
**
control
torso incision
esophagus inc.
head wedge
pharynx flank
48 hpi
D
#npp-1+ cells/1000µm²
*
control
pharynx flank
9dpi
E
pharynx area (cm)
worm length (mm)
r² = 0.8248

**Figure S4**

(**A**) Live images of injured animals immediately after injury, white arrows point to injury location. 200µm scale bar. (**B**) 20x confocal DAPI and H3P image representatives of each injury, 48hpi. 100µm scale bar. (**C**) Number of H3P+ cells in trunk region as defined by the white dotted box, H3P increase observed only for pharynx flank injury. one-way ANOVA with Dunnett's Test correction, n≥5.(**D**) Computer-aided, semi-automated counting of total *npp-1*+ cells per animal in each injury condition. (unpaired Student's t-test, n≥19). (**E**) Simple linear regression analysis of pharynx area versus fixed worm size, $r^2 = 0.8249$. (**C-D**) Symbols represent individual animals, data represented as box plot with mean +/- SD, *p<0.05, **p<0.01, ***p<0.001, ****p<0.0001, n.s. = p>0.05. Complete n-numbers in Table S1.

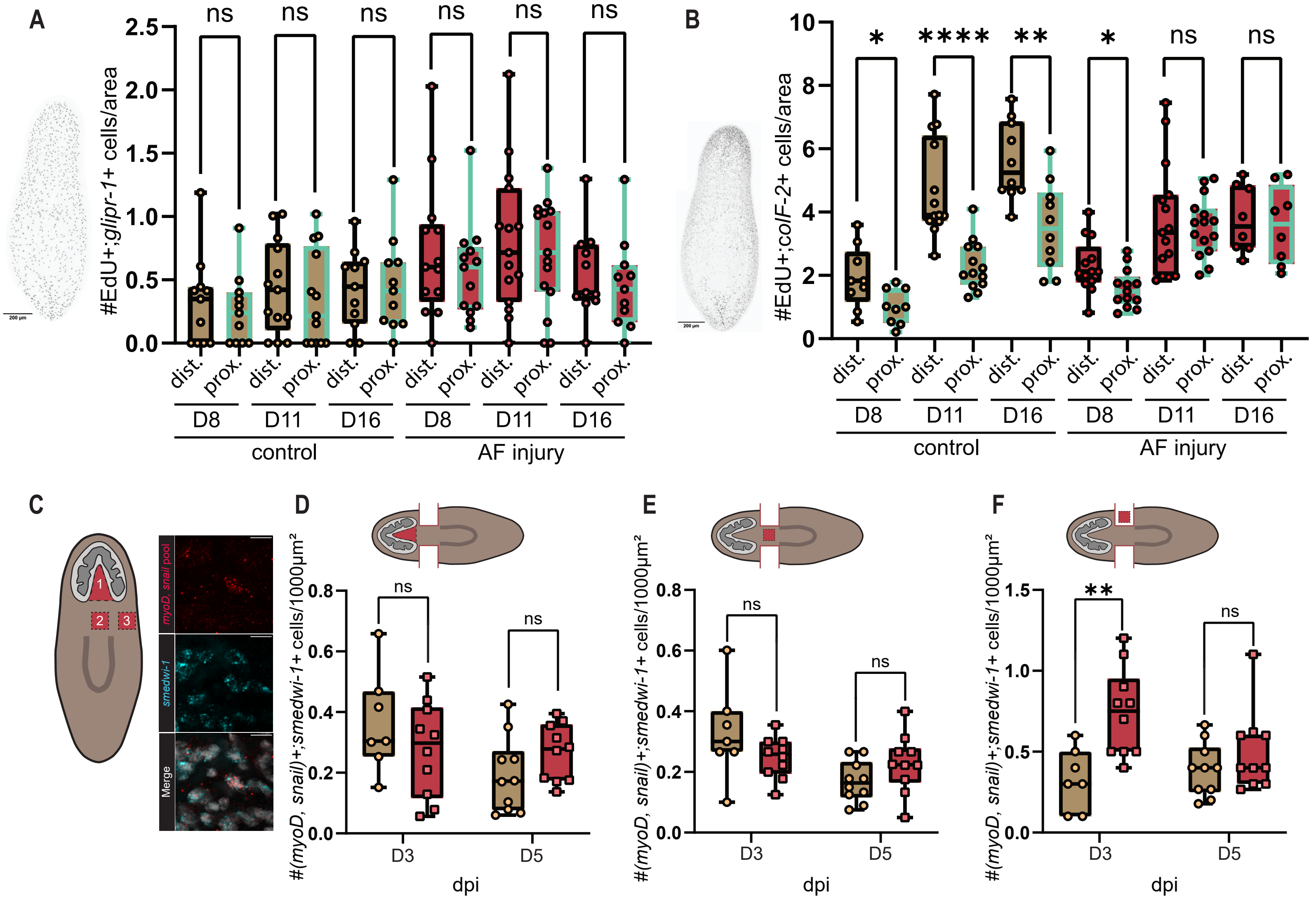

A
#EdU+;glipr-1+ cells/area
ns
dist.
prox.
D8
D11
D16
control
AF injury
B
#EdU+;colF-2+ cells/area
*
****
**
ns
C
1
2
3
myoD, snail pool
smedwi-1
Merge
D
E
F
#(myoD, snail)+;smedwi-1+ cells/1000µm²
D3
D5
dpi

**Figure S5**

(**A**) Number of EdU+; *glipr-1+* cells in wound-distal and wound-proximal areas per condition. Wound-distal is outlined in black, wound-proximal in cyan. Unpaired Student's t-test per condition, n≥11. (**B**) Number of EdU+; *colF-2+* cells in wound-distal and wound-proximal areas per (unpaired Student's t-test, n≥8). (**C**) Counting area schematic for L-N. *myoD+, snail+;smedwi-1+* cells. *myoD+, snail+;smedwi-1+* cells distal (n≥7) (**D**), proximal (n≥7) (**E**), in-blastema (n≥7) (**F**). (**A-B, D-F**) Symbols represent individual animals, data represented as box plot with mean +/- SD, *p<0.05, **p<0.01, ***p<0.001, ****p<0.0001, n.s. = p>0.05. Complete n-numbers in Table S1.

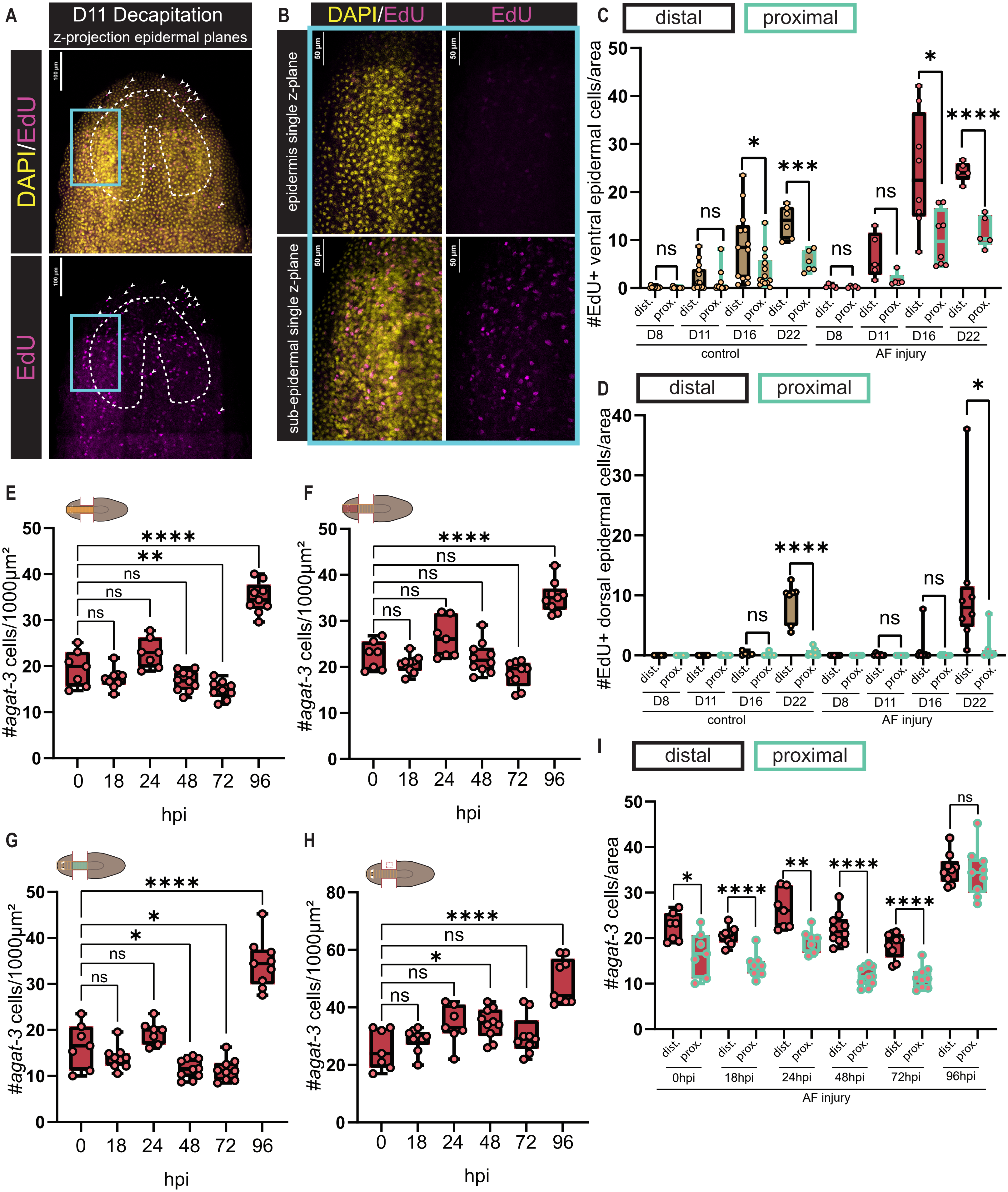
A
D11 Decapitation
z-projection epidermal planes
DAPI/EdU
EdU
B
DAPI/EdU
EdU
epidermis single z-plane
sub-epidermal single z-plane
C
distal
proximal
#EdU+ ventral epidermal cells/area
D8
D11
D16
D22
control
AF injury
D
#EdU+ dorsal epidermal cells/area
E
#agat-3 cells/1000μm²
hpi
F
G
H
I
#agat-3 cells/area
0hpi
18hpi
24hpi
48hpi
72hpi
96hpi

**Figure S6**

(A) The 40x single z-plane of epidermis in 11dpi decapitation animal shown in Fig. 6B. Bleed-through of signal from subepidermal DAPI is because of curvature of animal when imaging. 100µm scale bar. Blue rectangle indicates area of interest for B. (**B**) Top: zoomed in view of epidermal z-plane. Subepidermal plane of the same animal, showing the subepidermal DAPI that bleeds through into the epidermis plane. 50µm scale bars. (**C**) Number of EdU+ ventral epidermis cells in wound-distal and wound-proximal areas per condition. Black outlines indicate wound-distal, cyan indicate wound-proximal (unpaired Student's t-test, $n \geq 5$). **(D)** Number of EdU+ dorsal epidermis cells in wound-distal and wound-proximal areas per condition (unpaired Student's t-test, $n \geq 7$). (**E**) Computer-aided quantification of *agat-3+* progenitors in whole area, in distal area (**F**), in proximal area (**G),** and in-blastema (**H**), (one-way ANOVA with Dunnett's test correction). (**I**) Number of *agat-3+* cells in wound-distal and wound-proximal areas per condition. Black outlines indicate wound-distal, cyan indicate wound-proximal (unpaired Student's t-test). (**E-I**) $n \geq 7$. (**C-I**) Symbols represent individual animals, data represented as box plot with mean +/- SD, *$p<0.05$, **$p<0.01$, ***$p<0.001$, ****$p<0.0001$, n.s. = p>0.05n.s.: p>0.05. Complete n-numbers in Table S1.

**Table S1:** A list of n-numbers for all conditions in every figure panel.

**Table S2:** Summary table listing key details for each injury and experimental paradigm, including: the injury, whether it removed tissue, if it causes a generic wound response, if it causes a missing tissue response, time of fixation/analysis, EdU labeling relative to injury, length of EdU labeling, target tissue, proximity of injury to target tissue, and regenerative outcome.
